# Human-specific enrichment of schizophrenia risk-genes in callosal neurons of the developing neocortex

**DOI:** 10.1101/2021.09.10.459747

**Authors:** Emanuela Zuccaro, Vanessa Murek, Kwanho Kim, Hsu-Hsin Chen, Sara Mancinelli, Paul Oyler-Castrillo, Laura T. Jiménez-Barrón, Chiara Gerhardinger, Juliana R. Brown, Andrea Byrnes, Benjamin M. Neale, Joshua Z. Levin, Michael J. Ziller, Simona Lodato, Paola Arlotta

## Abstract

Human genetic studies have provided a wealth of information on genetic risk factors associated with neuropsychiatric diseases. However, whether different brain cell types are differentially affected in disease states and when in their development and maturation alterations occur is still poorly understood. Here we generated a longitudinal transcriptional map of excitatory projection neuron (PN) and inhibitory interneuron (IN) subtypes of the cerebral cortex, across a timeline of mouse embryonic and postnatal development, as well as fetal human cortex and human cortical organoids. We found that three types of gene signatures uniquely defined each cortical neuronal subtype: dynamic (developmental), adult (terminal), and constitutive (stable), with individual neuronal subtypes varying in the degree of similarity of their signatures between species. In particular, human callosal projection neurons (CPN) displayed the greatest species divergence, with molecular signatures highly enriched for non-coding, human-specific RNAs. Evaluating the association of neuronal class-specific signatures with neuropsychiatric disease risk genes using linkage disequilibrium score regression showed that schizophrenia risk genes were enriched in CPN identity signatures from human but not mouse cortex. Human cortical organoids confirmed the association with excitatory projection neurons. The data indicate that risk gene enrichment is both species- and cell type-specific. Our study reveals molecular determinants of cortical neuron diversification and identifies human callosal projection neurons as the most species-divergent population and a potentially vulnerable neuronal class in schizophrenia.

## Introduction

Large-scale human genetic studies of neurodevelopmental and neuropsychiatric disorders, such as Autism Spectrum Disorders [ASD], Bipolar Disorder [BD], and Schizophrenia [SCZ], have implicated hundreds of loci, including a variety of genes involved in neuronal development and function^1-10^. To date, the specific neuronal subtypes affected by these genetic variants are still poorly defined.

Comparing transcriptional profiles of neuronal subtypes with disease-enriched gene sets can potentially provide information on cell-type susceptibility; this approach has already begun to indicate links between individual disorders and cortical neurons, including associations of SCZ and BD with excitatory pyramidal neurons^11,12^. However, these experiments have to date largely relied on broadly-defined cell classes, with little subtype resolution. For example, most studies have not attempted to parse the diversity of cortical pyramidal excitatory cells, instead reporting broad pan-neuronal or pan-pyramidal associations. In addition, these experiments have largely examined adult cell types, or a limited set of developmental timepoints, which presents a limitation as the set of genes that are distinctive to a cell type and therefore important to its identity may change over development^13,14^. Finally, the gene sets specific for individual neuronal subtypes may differ between species^15^, thus requiring direct study of human neurons.

Here, we generated a high-resolution transcriptional map of multiple subtypes of both cortical excitatory projection neurons (PNs) and inhibitory interneurons (INs) over development, including embryonic and postnatal mouse neocortex, human fetal cortex, and human cortical organoids. We identify developmental, terminal, and stable molecular signatures for each of the major PN and IN subtypes in the mouse, and developmental signatures for the homologous human PNs classes. Notably, we find that expression of PN-subclass gene signatures diverges between the two species, with callosal projection neurons, the most evolutionarily recent PN class^16^, showing the greatest divergence. Leveraging these molecular signatures to identify neuronal types susceptible to genetic risk factors for various neuropsychiatric diseases, we show that polygenic risk for SCZ is significantly enriched in signature genes of the CPN class in human fetal cortex. This enrichment was specific to human CPNs, suggesting that SCZ risk genes may have species-specific effects on neurodevelopment.

## Results

### Molecular signatures with distinct temporal dynamics collectively define pyramidal neuron and interneuron subtype identity in the neocortex

In order to trace the dynamic molecular signatures of individual neuron classes, we profiled the major subtypes of cortical excitatory and inhibitory neurons at six different time points along a timeline spanning phases of neuronal development from early postmitotic fate decisions and neuronal migration to synaptic integration and circuit maturation.

We first transcriptionally profiled the three major subtypes of excitatory PNs, corticothalamic PNs (CThPNs) of layer 6, subcerebral PNs (ScPNs) of layer 5, and CPNs of both deep and upper layers, in the mouse cortex (Figure 1A and Table S1). To simultaneously and systematically purify multiple PN subtypes from the same sample, we employed an optimized version of the MARIS technique^17,18^, where the combinatorial expression of three transcription factors, BCL11B, TLE4, and SATB2^19-22^, is used to isolate the three major cortical PN subtypes. Although this approach requires pre-defining the target cell populations, it allows deeper sequencing compared to single-cell RNA sequencing. We FACS-purified ScPNs as BCL11B^high^/TLE4^low^/SATB2^-^, CThPNs as BCL11B^low^/TLE4^high^/SATB2^-^, and CPNs as BCL11B^-^/TLE4^-^/SATB2^+^ from the mouse somatosensory cortex at six different time points (E16.5, E18.5, P1, P3, P7, and P30, n≥3 per stage, 48 FACS-purified and transcriptionally profiled samples) (Figure 1A and Figure S1A-C) (Table S1).

**Figure 1:**
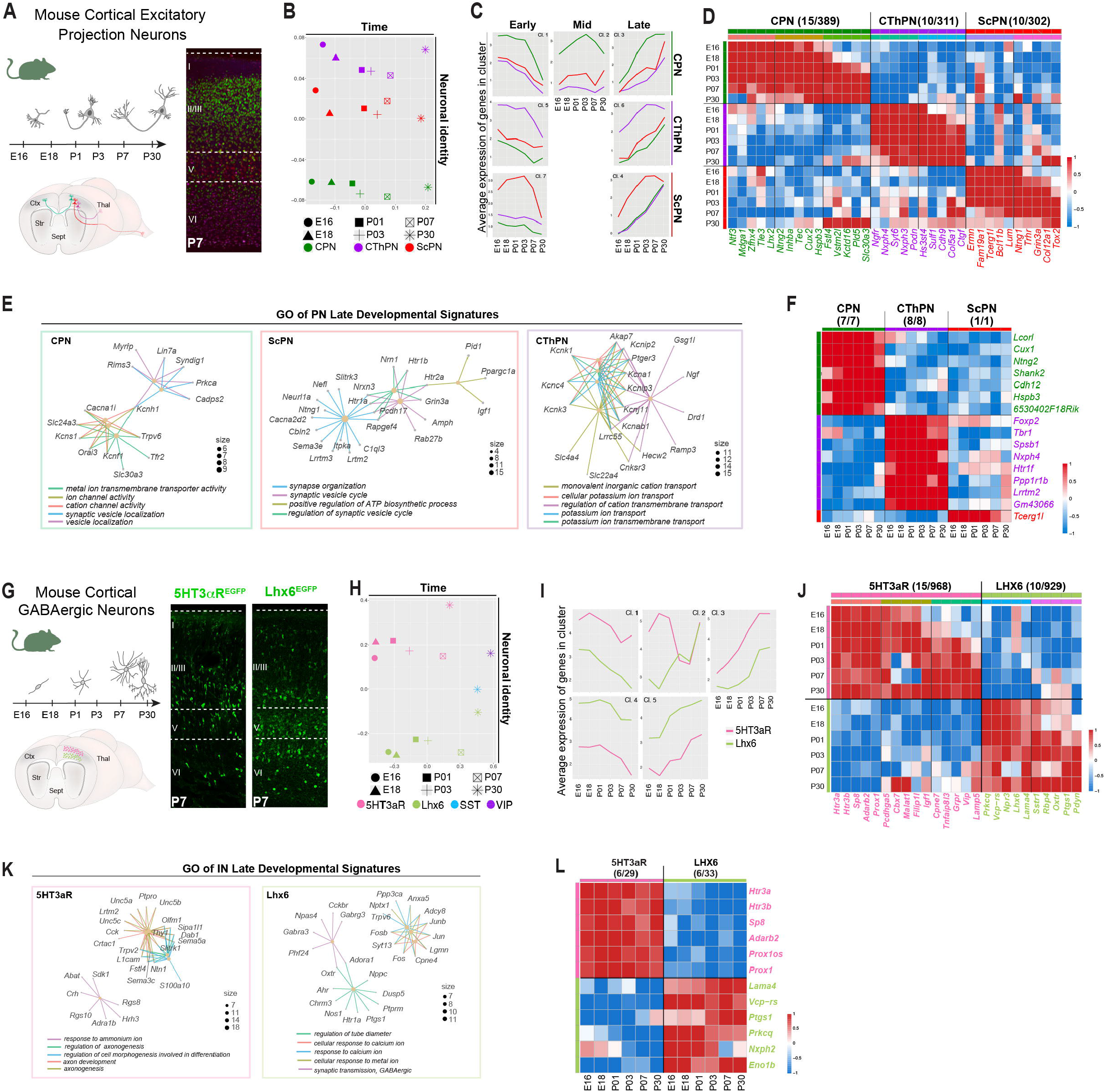
FACS-Purification and transcriptional profiling of mouse cortical PN and IN subtypes along embryonic and postnatal development. **A**) Schematic diagram of samples collection. CPNs (green), ScPNs (red), and CThPNs (violet) were simultaneously isolated from dissected somatosensory cortices across six developmental stages from E16 to P30. Representative images of immunofluorescence for SATB2 (CPN marker, green), BCL11B (ScPN marker, red), and TLE4 (CThPN marker, violet) on mouse cerebral cortex sections at P7. Details on biological replicates, library sample size, quality and RNA sequencing parameters are reported in Table S1. **B**) Dimensionality reduction by multidimensional scaling (MDS) of the average gene expression shows that PN sample dissimilarity is primarily driven by time (Dimension 1) and identity (Dimension 2). **C)** Line plots of average gene expression illustrating cluster association with distinct subtype identities and developmental stages. PN subtype-specific developmental signatures - obtained as explained in detail in the Methods - were classified in early, mid, and late, reflecting the time-specific expression dynamic along development. Differentially expressed signature genes of PN subtypes are reported in Table S2. Clusters 1, 5, and 7 identify early developmental signatures for CPNs, CThPNs and ScPNs, respectively; cluster 2 represents CPN mid-developmental signature genes; clusters 3, 6, and 4 represent CPN-, CThPN, and ScPN-late developmental signatures, respectively. Gene clusters are reported in Table S3. **D**) Heatmap showing expression pattern of PN subtype-specific developmental signature genes (5 top-ranked/cluster genes are shown). **E)** Cnet plots displaying gene concept network analysis for GO terms enriched in CPN- (green square), ScPN- (red square), and CThPN- (violet square) subtype specific late-developmental signatures. Gene Ontology (GO) analysis reveals enrichment of genes associated with circuit establishment and maintenance. Statistical details and full description of GO terms is reported in Table S4. **F)** z-scores expression heatmap of PN subtype-specific stable signatures. **G**) Schematic overview of purification strategy of 5HT3aR- (pink) and Lhx6- (light green) expressing cortical INs. Representative images of P7 cerebral cortex from genetically labeled 5HT3aR-GFP and Lhx6-EGFP mouse IN reporter lines. **H**) Dimensionality reduction by multidimensional scaling (MDS) of average gene expression shows that IN sample dissimilarity is primarily driven by time (Dimension 1) and class identity (Dimension 2). **I)** Line plots of average gene expression illustrating cluster association with distinct subtype identities and developmental stages. Early, mid, and late developmental signatures reflected the expression dynamics along development. IN subtype-specific developmental signatures - obtained as explained in detail in the Methods - were classified in early, mid, and late, reflecting the time-specific expression dynamic along development. Differentially expressed signature genes of IN subtypes are reported in Table S2. Clusters 1 and 4 identify early-development signatures for 5HT3aR and Lhx6, respectively; cluster 2 represents 5HT3aR mid-developmental signature genes; clusters 3 and 5 represent late developmental signatures of 5HT3aR and Lhx6, respectively. Gene clusters are reported in Table S3. **J**) Heatmap showing expression pattern of IN subtype-specific developmental signature genes (5 top-ranked/cluster genes are shown). **K**) Cnet plots displaying gene concept network analysis for GO terms enriched in 5HT3aR- (pink square) and Lhx6- (light green square) subtype specific late-developmental signatures. Enrichment of genes associated with axonal development (5HT3aR) and synaptic transmission (Lhx6) were found. Statistical details and full description of GO terms is reported in Table S4. **L**) z-scores expression heatmap of IN subtype-specific stable signatures. Abbreviations: E, embryonic day; P, postnatal day; CPN, Callosal Projection Neurons; ScPN, Subcerebral Projection Neurons; CThPN, CorticoThalamic Projection Neurons; IN, Interneurons; SST, Somatostatin; VIP, Vasointestinal peptide. Scale bars: 250μm.

To begin to explore the transcriptional and temporal dynamics defining PN diversity, we visualized the overall variation in the dataset using dimensionality reduction by Multidimensional Scaling (MDS) on average gene expression between the PN subtypes along their development. This showed that sample dissimilarity was primarily driven by time (Dimension 1) and neuronal identity (Dimension 2) (Figure 1B). Consistent with their known expression patterns *in vivo*, we observed increased expression of *Bcl11b, Tle4*, and *Satb2* in ScPNs, CThPNs, and CPNs, respectively, while markers of other cell types (e.g., oligodendrocytes, endothelial cells, interneurons, astrocytes) showed negligible expression (31 out of 39 genes had FPKM < 2.5), thus validating the specificity of our FACS-purification strategy (Figure S1G-L).

We considered both time and neuronal subtype identity to determine differentially-expressed genes between all pairwise comparison (DEGs; Figure S1d-f, see Methods for criteria). We then analyzed the resulting dataset to extract *developmental, stable*, and *terminal* molecular signatures uniquely identifying ScPNs, CThPNs, and CPNs (Figure 1C-D,F and Figure S4A). To discover developmental signatures, we applied stringent criteria to filter out transcriptional changes reflecting the asynchronous maturation of cortical PNs (see Methods), retaining transcripts whose level of expression was enriched in only one subtype for at least two consecutive time points (1,008 genes, Table S2). Clustering the resulting set using partitioning around medoids (PAM, k-medoids) identified seven clusters of genes with distinct temporal and subtype-specific expression patterns, corresponding to early, mid, and late stages of development of each neuronal class (Figure 1C-D and Figure S2A). These clusters contained both established (e.g, *Ntng2* and *Inhba* for CPNs; *Wnt7b* and *Kcnab1* for CThPNs; *Lum, Pex5l*, and *Grik2* for ScPNs)^23-25^, and novel (e.g., *Fstl4, Nr1d2, Cadps2* for CPNs; *Ssrt2, Lig2* and *Fap* for CThPNs; *Tox2, Col12a1*, for ScPNs) subtype-specific markers (Figure 1D and Figure S2A and Table S3). Gene ontology analysis of the developmental signatures for each PN class revealed expected enrichment of developmental processes such as axonogenesis in early-to mid-developmental signatures, and synaptic maturation and function in late-developmental signatures (Figure 1E, Figure S2B, and Table S4).

We next examined this gene set to identify *stable* signatures for PN subtypes, i.e., genes consistently enriched in a single PN class at all time points, regardless of developmental stage. For both CPNs and CThPNs, stable signature genes included novel coding transcripts and unmapped loci (*Lcorl, Shank2 a*nd *6530402F18Rik*), as well as canonical markers already known to be involved in PN development (i.e., *Cux1, Hspb3, Foxp2, Tbr1*)^18,26^ (Figure 1F, Figure S2C, and Table S5).

Finally, we identified *terminal* signatures, which we strictly defined as subtype-specific transcripts whose enrichment emerged exclusively at P30, after definitive transcriptional identity is established, and in only one neuronal subtype (Figure S4A and Table S5). We identified 136 subtype-specific terminal signature genes, which included genes associated with processes such as dendritic spine development and plasticity (e.g., *Baiap2l2* in ScPNs)^27^, synaptic connection and trans-synaptic signaling (e.g., *Teneurin* and *Tcap* in CPNs), and neuro-immune response (e.g., *Il11ra1* in CThPNs) (Figure S4A and Table S4). The subtype-specific expression pattern of these genes was consistent with single-cell resolution data from mature cortical subtypes in the Allen Mouse Cell Types Database^28^ (Figure S4C). These results show that the molecular class identity of terminally-differentiated neurons is defined largely by the genes used to execute their functional properties.

All PN types wire into a local cortical microcircuitry with distinct classes of cortical inhibitory interneurons (INs)^29^. Because INs display a high degree of cellular, molecular and functional diversity^30,31^, we sought to also define the molecular signatures of cortical IN populations through time. Cortical interneurons derive from two main germinal zones, the medial ganglionic eminence (MGE), and the caudal ganglionic eminence (CGE) ^30^. We used genetically-labeled mouse lines to isolate MGE-derived (Lhx6-GFP) and CGE-derived (5Ht3aR-GFP) cortical INs from the somatosensory cortex at the same developmental stages used for PNs, as well as SST-dtTomato and VIP-tdTomato reporter lines to isolate mature somatostatin (SST) and vasoactive intestinal polypeptide (VIP) IN subtypes at P30 (n≥3 per stage, Table S1, 35 libraries, Figure 1G). As for PNs, dimensionality reduction using MDS indicated that sample dissimilarity was primarily driven by time (Dimension 1) and neuronal type (Dimension 2) (Figure 1H).

From this dataset, we performed pairwise differential gene expression analysis across all developmental stages (filtering criteria in Methods, Figure S3a, see Methods) and extracted *developmental, terminal*, and *stable* signatures using the same approach described for PNs (Figure 1I-J,L, Figure S3C, and Table S2-3,5). Divergence between the two cardinal IN subclasses (Lhx6- and 5Ht3aR-lineage) was evident from the earliest stages of development (Figure 1G, Figure S3B-C, and see Methods). In contrast to PNs, however, IN developmental signatures (1,789 genes, Table S2) included a large fraction of genes stably enriched in one population alone (Figure 1L and Figure S3E), confirming that the two cardinal subdivisions (5Ht3aR and LHX6) follow largely non-overlapping identity programs from the earliest stages of development. GO analysis revealed processes appropriate for each developmental stage for both IN subdivisions. Interestingly, late signature genes of Lhx6-positive INs displayed specific enrichment for genes involved in synaptic transmission, while signatures of 5HT3aR-positive INs were enriched for axonogenesis genes (Table S4).

Terminal signatures of interneurons emerged at later stages, by P30, when distinctive molecular features of each IN subtype have appeared (such as *Crhb4, Htr1a*, and *Oprm1* for SST-INs and *Tac2, Frem1*, and *Npy1r* for VIP-INs). These remained enriched in the adult cortex (Figure S4B, D, Allen Mouse Cell Types Database^28^ and Table S5). This observation supports a late specification of IN subtypes, likely following recognition and pairing with their PN partners and integration into the cortical local circuit.

All together, these data indicate that transcriptional signatures that define individual neuron class identity in the neocortex are mostly composed of genes that change expression over development, rather than invariant signatures. The fact that cell identity signatures are mostly dynamic indicates that no single time point exemplifies a cell type, highlighting the need to include more complete timelines of development and maturation to comprehensively capture signatures of cell identity.

### Molecular diversity of ScPNs in distinct functional areas

The cerebral cortex is divided into multiple functional areas^32,33^. To begin to examine whether the signatures we identified might describe individual PN types across multiple regions of the cortex, we took ScPNs of layer Vb as a test case and compared them across areas. For ScPNs, the only stable signature gene (i.e., differentially expressed at all ages) was *Tcerg1l*, a previously identified ScPN marker^18,25^. We therefore employed a tamoxifen-inducible *Tcerg1l*-CRE driver mouse line^34^ crossed to a Sun1:GFP nuclear reporter^35^ to label and isolate mature layer V ScPNs across different functional areas. We FACS-isolated and profiled *Tcerg1l* lineage nuclei from P56 *Tcerg1l*/Sun1:GFP mice (n ≥ 4 replicates per cortical region, each representing a pool of 5-6 mice) from motor (M), somatosensory (SS), auditory (AUD) and visual cortex (VIS), each sampled at 1 or 2 defined anterior-posterior (AP) levels, and performed single-nucleus RNA sequencing (snRNA-seq) (Figure 2 A-B, Figure S5).

**Figure 2:**
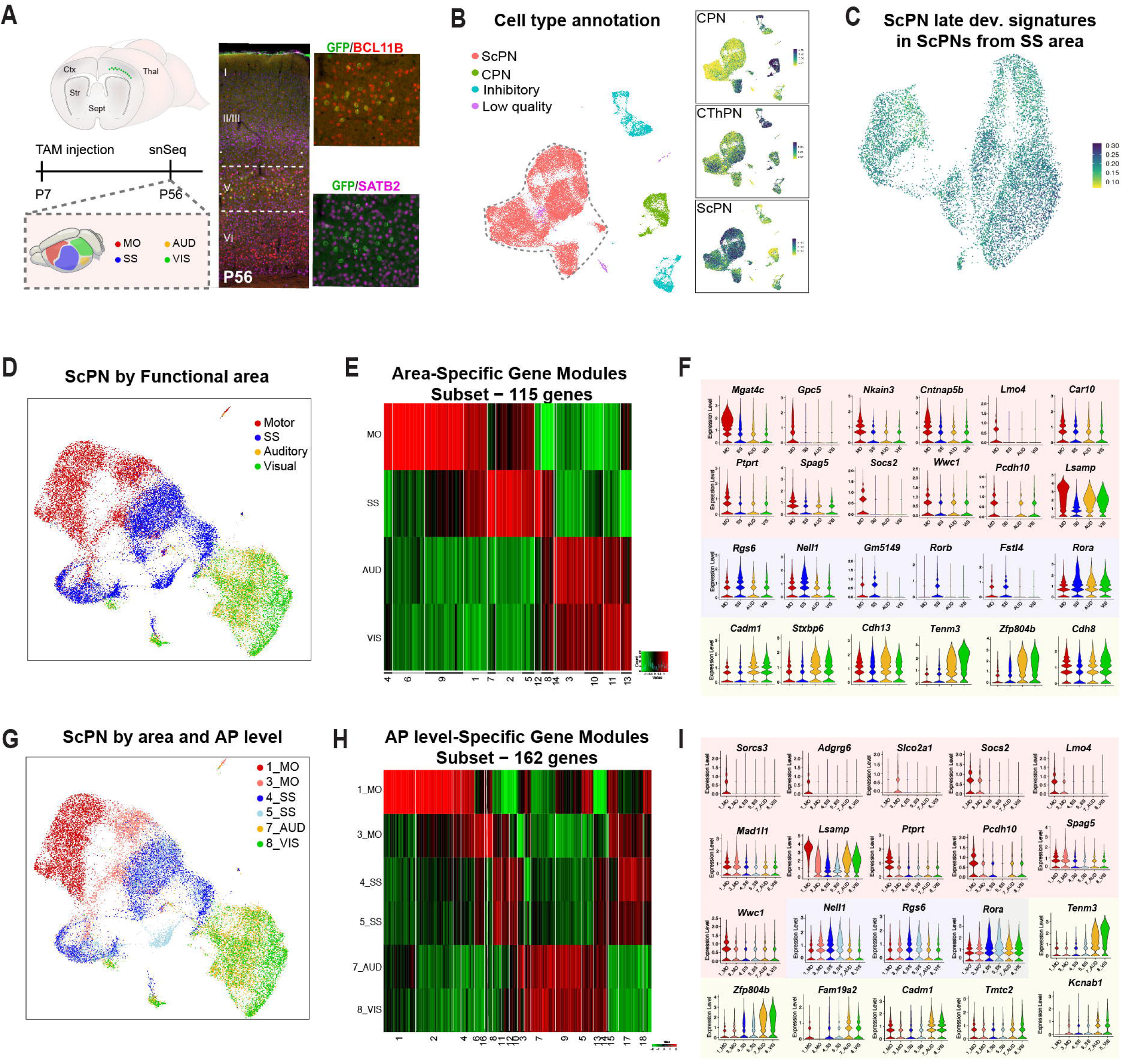
ScPN molecular diversity across functional areas and along the anterior-posterior axis. **A)** Schematic diagram of experimental design. *Tcerg1l*-CreERT2/Sun1:GFP mice were injected with Tamoxifen at P7 and corteces analyzed at P56. Representative images of double immunofluorescence of BCL11B (red) and SATB2 (violet) showing colocalization with GFP-expressing cells (proxy for *Tcerg1l*-CreERT2/Sun1:GFP). Scale bars: 250μm. Zoom-in inset: scale bar: 100μm. **B**) Uniform Manifold Approximation and Projection (UMAP) plots of Tcerg1l-GFP expressing nuclei isolated by FACS confirming that the majority of the recovered nuclei in all the areas are ScPNs, while a smaller fraction also contains CPNs and CthPNs, as well as INs. **C**) UMAP of computationally extracted ScPNs from B) (only SS area), showing a homogenous enrichment of late ScPN signature genes among nuclei isolated from somatosensory cortex. **D**) UMAP of ScPNs in distinct functional areas of the cerebral cortex. **E**) Heatmap of area-specific expression modules derived from differential gene expression analysis of ScPNs between cortical functional areas. **F)** Representative violin plots of ScPN signature genes of motor (red), somatosensory (blue), visual (green) and auditory (yellow) areas. Differentially expressed signature genes of ScPN across areas are reported in Table S6. **G**) UMAP of ScPNs in distinct anterior-posterior levels of different cortical areas. **H**) Heatmap of AP level-specific expression modules derived from differential gene expression analysis of ScPNs between distinct rostro-caudal locations. **I)** Representative violin plots of ScPN signature genes of motor (red/pink), somatosensory (blue/light blue), visual (green) and auditory (yellow) areas. Differentially expressed signature genes of ScPN across AP levels are reported in Table S6. Abbreviations: Mo, Motor; SS, Somatosensory; AUS, Auditory; VIS, Visual; CPN, Callosal Projection Neurons; ScPN, Subcerebral Projection Neurons; CThPN, CorticoThalamic Projection Neurons; AP, anterior-posterior.

We first confirmed that the sorted nuclei expressed the ScPN late-developmental signature; as expected, the majority of the nuclei recovered expressed this signature (Figure 2B and Figure S5D). This line also labels a small number of CPNs and GABAergic interneurons, which we correctly detected in our dataset (*RorB+*-CPNs, 8.8%, and GABAergic interneurons, 11.7%); these cells were excluded from further analysis (Figure S5D).

We then assessed the degree of consistency of the ScPN late-developmental signature among all ScPN nuclei isolated from the primary somatosensory cortical area, and found that it was comparable across all nuclei (Figure 2C, Figure S5E). Moreover, this signature was equivalently expressed at different AP levels and in all functional primary areas sampled (Figure S5F), indicating that it represents a core molecular feature of ScPN identity across multiple areas of the mature brain.

In addition to this shared signature, we found that each functional area displayed a unique molecular identity, with ScPN nuclei clustering according to their anatomical location (Figure 2J-M and Figure S5H). We found 14 gene modules (see Methods) that varied between ScPNs of different cortical functional areas (e.g., motor cortex ScPNs differentially express *Lmo4, Socs2, Mgat4c*, and *Gpc5*), and 18 gene modules that varied over both functional area and AP position (Figure 2L and Table S6). The data highlights an additional level of molecular heterogeneity of the ScPN population in the adult brain, both between different areas and between different AP positions within the same area.

The findings indicate that the core molecular programs that define the late-developmental stages of neuronal subtype-specific identity continue to be broadly active in the adult cortex, but that area-specific signatures are also present.

### Cross-species comparison reveals that human callosal projection neurons display the greatest transcriptional divergence

The cerebral cortex has diverged substantially during primate evolution, including molecular differences between corresponding cell types in different species^15,36,37^. Although a number of reports have profiled adult human pyramidal neurons to disentangle their transcriptional and epigenetic heterogeneity^38-40^, the molecular signatures of developing human PN subtypes are poorly defined. These are needed to investigate linkage to human disease, to allow cross-species comparative analysis, and for validation of the class identity of PN subtypes generated *in vitro*^41^.

We therefore sought to define the molecular identity of PN types from the human fetal cortex. We applied the same experimental approach used for murine tissues to FACS-isolate and transcriptionally profile three molecularly distinct excitatory populations from the human developing cortex at six gestational stages (gestational week [GW]16, 17, 18, 19, 20 and 21, n≥2 per stage; Figure 3A-B, Figure S6A and Table S1), based on the subtype-specific expression of the three TFs *BCL11B, TLE4*, and *SATB2* (Figure 3C-D; Figure S6B-C, and Table S1). This dataset embodies ground-truth information on pre-defined cortical PN diversity through human mid-gestation.

**Figure 3:**
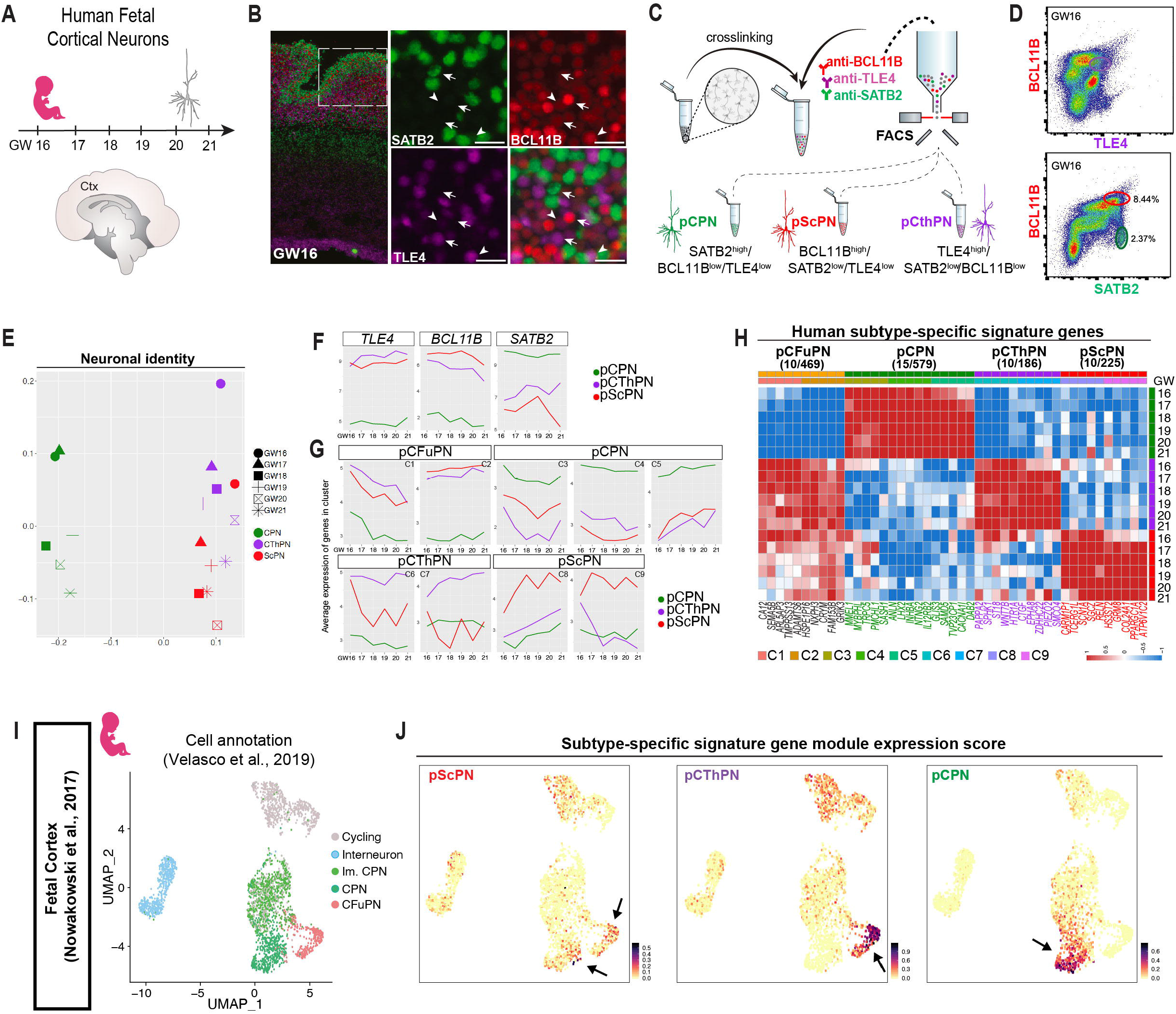
Identification, FACS-isolation, and transcriptional profiling of PN subtypes from human fetal cortices at midgestation. **A**) Schematic of sample collection. **B**) Representative images of human fetal cortical sections isolated from GW16 and immunostained for SATB2 (labeling pCPNs in green), BCL11B (marker of ScPNs in red), and TLE4 (labeling CThPNs in violet). Arrows indicate pScPNs expressing high levels of BCL11B and low levels of SATB2, while arrowheads represent pCThPNs expressing high levels of TLE4 and low levels of BCL11B. Scale bars: 250μm. Zoom-in inset: scale bar: 100μm. **C**) FACS-isolation gating strategy of molecularly identified human cortical populations. **D**) Representative FACS plots identifying pCPNs (BCL11B^-^ /TLE4^-^/SATB2^+^), pScPNs (BCL11B^high^/TLE4^low^/SATB2^-^), and pCThPNs (BCL11B^low^/TLE4^high^/SATB2^-^). Details on biological replicates, library sample size, quality and RNA sequencing parameters are reported in Table S1. **E**) Dimensionality reduction by multidimensional scaling (MDS) of average gene expression shows that the samples dissimilarity is driven by identity in the first two dimensions, clearly separating pCPNs (represented in green) from pCFuPN in the first dimension and to a lesser degree CThPN (represented in violet) and pScPN (represented in red) in the first dimension. The second dimension shows a less evident trend in sample separation by time. **F**) Line plots representing the expression profile of *Tle4, Satb2*, and *Bcl11b* transcripts. Gene expression is in accordance with sample identity and consistent over time. **G**) Line plots of average gene expression illustrating cluster association with distinct subtype identities and developmental stages. PN subtype-specific signatures - obtained as explained in detail in the Methods - are reported in Table S2. Clusters 1 and 2 represent pCFuPN signatures, with genes highly expressed in both pScPN (red line) and pCThPN (violet line), while depleted in pCPN (green line); Clusters 3, 4, and 5 were representing genes enriched in pCPN; Clusters 6 and 7 pCThPN genes; and Clusters 8 and 9 genes enriched in pScPN. Gene clusters are reported in Table S3. **H**) z-score heatmap representing the expression pattern of human subtype-specific developmental signature genes at different ages (5 top-ranked/cluster genes are shown). **I**) UMAP of a subset of human fetal cortex cells from the previously published single cell study^43^. **J**) Re-annotation of dataset in (**I**) using our subtype-specific signatures (identified in **G**). Subpopulation of pScPNs and pCThPNs, previously annotated as “CFuPN”, are now clearly mapped and indicated with arrows. Abbreviations: GW, gestational week; pCPN, putative Callosal Projection Neurons; pScPN, putative Subcerebral Projection Neurons; pCThPN, putative CorticoThalamic Projection Neurons; pCFuPNs, putative Corticofugal Projection Neurons; Im CPN, immature CPN.

Dataset visualization using MDS showed that during the sampled developmental window, the three neuronal populations clustered based on their identity, with CPNs (*SATB2*^high^/*BCL11B*^low^/*TLE4*^low^) distinctly segregated from the two corticofugal (CFuPNs) populations (Figure 3E), and CThPNs (*TLE4*^high^/*BCL11B*^low^/*SATB2*^-^) separated from ScPNs (*BCL11B*^high^/*TLE4*^low^/*SATB2*^-^), although they were less resolved from each other than their murine counterparts. We confirmed the expression of genes known to be PN subtype-specific, while expression of glial, oligodendrocyte and non-neuronal markers were virtually absent, validating our isolation approach (Figure S6E-J). Of note, contrary to the mouse neurons, human samples did not show a clear separation based on gestational age, suggesting that, consistent with the prolonged length of human brain development, the selected six-week time window may only cover limited developmental progression within each neuronal subtype (Figure 3E).

We identified subtype-specific signatures using criteria similar to those used to define developmental signatures in our mouse analysis, omitting the requirement for specific enrichment in two consecutive time points, as time was not a major driver of variation in this dataset. This identified 2,299 DEG; PAM clustering identified multiple gene clusters with distinct subtype expression patterns, including clusters specific for putative (p) pCThPNs, pScPNs, pCPNs, and corticofugal projection neurons (pCFuPNs) (Figure 3F,H) (Table S7). These signatures contained both known markers of the homologous murine PN subtypes (e.g., *CUX2* and *LHX2* for pCPNs, *NFIA* and *PAPPA2* for pCThPNs, and *TCERG1L* and *S100A10* for pScPNs), as well as genes with human-specific expression (Figure 3H), which we validated using two single cell transcriptomic datasets of human fetal cortex^42,43^ (Figure 3I and Figure S7A-B, see Methods). Using our gene signatures, we were able to identify pScPN and pCThPN populations in these datasets, as well as the previously annotated upper-layer PNs (Figure 3L and Figure S7A). For all PN subtypes, a subset of the signature genes identified in our fetal dataset also retained subtype-specific expression in the adult human cortex, as assessed by *in situ* hybridization (Figure S8A)^44^ and spatial transcriptomics of human dorsolateral pre-frontal cortex (DPFC)^45^ (Figure S8B) (*LPL* and *NTNG2* for CPNs, *HS3ST2* and *COL24a1* for ScPNs, and *PP1R1B* and *ST18* for CThPNs, respectively).

These data indicate that during mid-gestation, human PN subtype signatures already comprise genes that define their adult identity. Although human late-fetal and postnatal molecular PN development is yet to be mapped, the persistent subtype-specific expression of fetal signature genes in the adult suggests the existence of stable signatures for human cortical PN classes.

To investigate the phylogenic conservation of PN molecular identity between mouse and human, we next compared their respective PN signature gene sets. We identified 1803 genes with a corresponding ortholog in the other species, while 577 genes in the human signatures had no mouse ortholog, mainly non-coding RNAs or annotated pseudogenes (Table S8). Notably, 60.3% of those genes belonged to the human CPN signatures, while 39.7% were found in the ScPN and CThPN signatures. The human CPN signature was correspondingly the most enriched for human-specific genes (56% of CPN signature genes, 43% of ScPN, 43% of CThPN, 31% of CFuPN).

Extending this analysis to other members of the primate clade (including Apes, Old World Monkeys, New World Monkeys, and Promisians) showed that over 90% of the genes not shared with mouse (i.e., *Nlgn4x, Diras3* and *Zim2*) were not conserved in other primates and are exclusively present in the human genome (Figure 4A). CPNs evolved relatively recently compared to the other cortical PN subtypes and have undergone a disproportionately large population expansion from mouse to human^16^. Our data suggest that many of the genes that define the molecular identity of this neuronal class in humans are of an evolutionarily recent origin.

**Figure 4:**
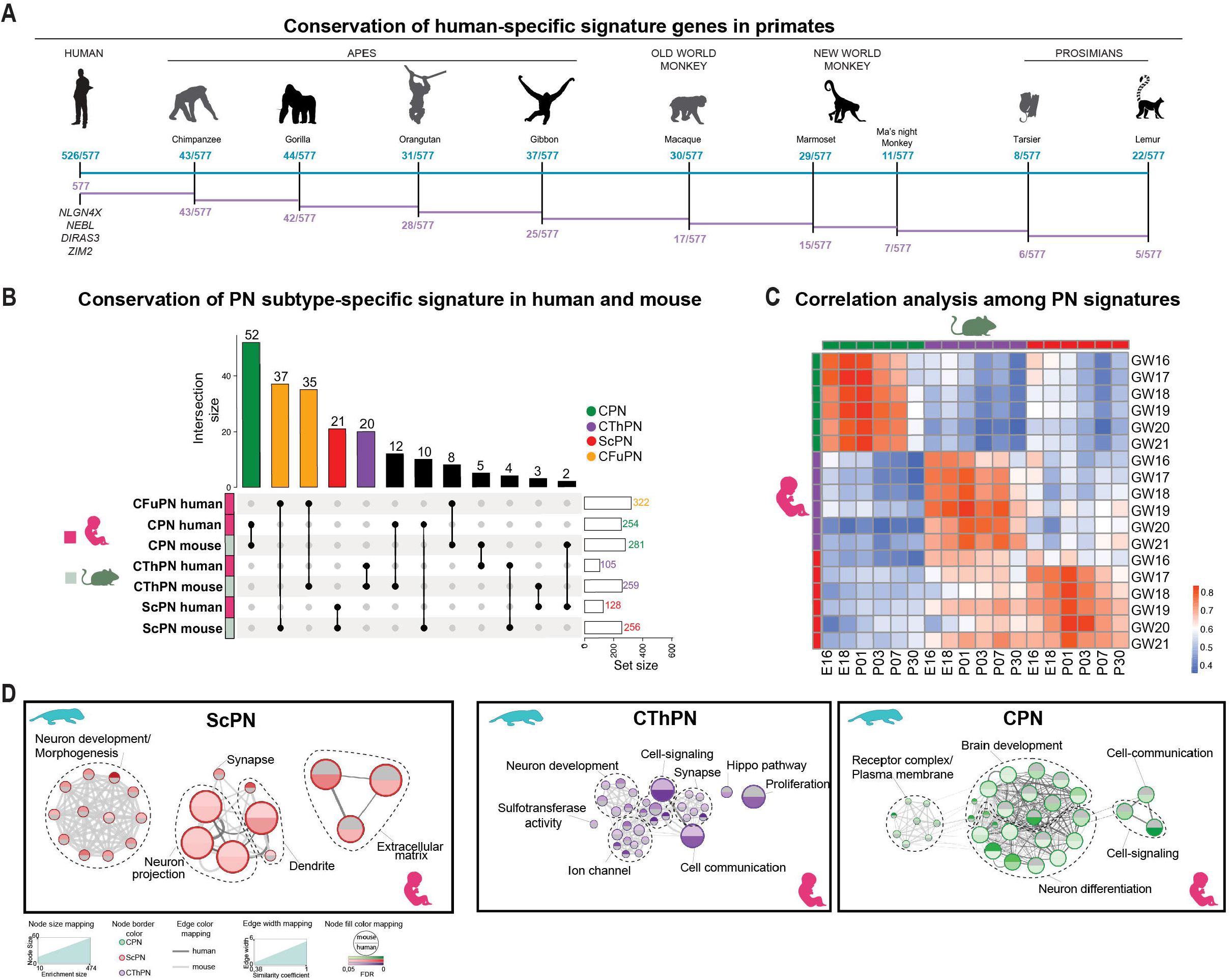
Interspecies comparison of human and mouse subtype-specific PN signature genes. **A**) Occurrence of the human-specific (no ortholog in mouse) signature genes in the various primate clades. Assignment of the 577 human-specific signature genes (Table S8) to a primate clade, based on the primate genome(s) in which an ortholog was found in the present analysis (BioMart database). Clades are specified in top panel. The color-coding indicates the degree of conservation among species in each clade: the number reported in blue represents the frequency of genes that are present both in human and the given primate species; the number in violet represents the frequency of genes present in multiple primate genomes within the prosimian and simian groups. **B**) Upset plot representing gene set intersection between human and mouse ortholog subtype-specific signature genes. Only a subset of identified signature genes for projection neuron populations intersect, while the majority of projection neuron signature genes do not intersect. **C**) Pearson’s correlation of gene expression of subtype-specific ortholog signatures in human and mouse at the different sampled ages. Highest correlation between samples of corresponding subtype across species was observed at P1. **D**) G-profiler based enrichment analysis of human signature and mouse DEG at P1, and network visualization of shared and species-specific enriched terms by Cytoscape analysis (see Methods). Statistical details and enrichment term description is reported in Table S9. Abbreviations: GW, gestational week; E, embryonic day; pCPN, putative Callosal Projection Neurons; pScPN, putative Subcerebral Projection Neurons; pCThPN, putative CorticoThalamic Projection Neurons; pCFuPNs, putative Corticofugal Projection Neurons; Im CPN, immature CPN; FDR, false discovery rate.

Next, we systematically performed cross-species comparison between transcriptional signatures of mouse and human cortical PNs during development (Figure 4 B-C and Figure S9A). We first applied MDS on all PN samples from both species using the genes from the orthologous signature gene lists, which revealed sample separation based primarily on molecular identity (Figure S9A). To address the degree of homology of the signature sets and identify conserved and human-specific developmental pathways, we directly quantified the percentage of developmental signature genes shared between the two species. For each human subtype, approximately 20% of signature genes were shared with the mouse counterpart (20.5% for CPN, 16.4% for ScPN, 19% for CThPN), while 7-18% of the mouse developmental signature genes were shared with the corresponding human neuronal subtype (18% for CPN, 7% for ScPN, 8% for CThPN), suggesting a limited conservation of subtype-specific gene programs between the two species (Figure 4B).

To assess whether this low overlap might stem from differences in the developmental stage of the human samples compared to the mouse, we computed Pearson correlation coefficients on the orthologous genes. This analysis showed that human PN subtypes isolated from midgestation cortex (GW16-21) are transcriptionally more similar to PN subtypes from P1 mouse cortex (Figure 4C). The correlation to the P1 mouse samples was consistent across all PN classes, indicating that, although human fetal cortex development spans a longer period than in mice, it also occurs earlier relative to birth. This may reflect molecular events which unfold postnatally in mice but occur during fetal stages in human cortical development.

Given this specific correlation between human midgestation and the murine P1 stage, we focused on this stage for further comparative analysis. We sought to investigate the degree of overlap of cellular features, molecular pathways, and biological processes shared among the two species at this specific stage, by employing an established pipeline for pathway enrichment (PE) analysis^46^. We examined the genes with orthologs in both species and determined P1-enriched DEGs for each PN type (Table S8, see Methods). PE analysis on these genes identified a dense core of nodes (enriched gene ontology terms) and edges (common genes among nodes) in support of shared pathways underlying multiple developmental processes (Figure 4D). This indicates that human and mouse PNs rely on different genes (as shown by their divergent signatures) to execute similar biological functions at this stage. However, our approach also revealed species-specific enrichment for some terms: cell communication for human CPN signatures and extracellular matrix for human ScPN signatures (Table S9, Figure 4D). This suggests the formation of a cellular environment unique to human corticogenesis.

### Human fetal CPNs display a specific enrichment for SCZ risk genes

To determine whether the subtype-specific molecular maps we built for human and mouse cortical development could be used to assess disease susceptibility in distinct neuronal subtypes and along multiple developmental stages, we employed linkage disequilibrium (LD) score regression^47^ in combination with a large panel of GWAS summary statistics for neuropsychiatric diseases and other complex diseases^48^. Several non-neuropsychiatric traits were included as controls (Table S10). For each of our mouse developmental and human fetal signature gene sets, we identified a window of 100kb upstream and downstream of the canonical transcriptional start site (TSS) for each gene, and calculated risk association via prior heritability enrichment, as previously described^52^. For mouse, we analyzed the corresponding human orthologs (Table S8).

For the mouse datasets, this analysis showed no enrichment in any signature gene set of either PN or IN subclasses, for any of the traits examined. To verify whether individual associations existed when subpopulations of mouse PNs and INs were analyzed individually, we next clustered each of them separately. Again, we did not observe any significant associations. (Figure 5A-B and Figure S10).

**Figure 5:**
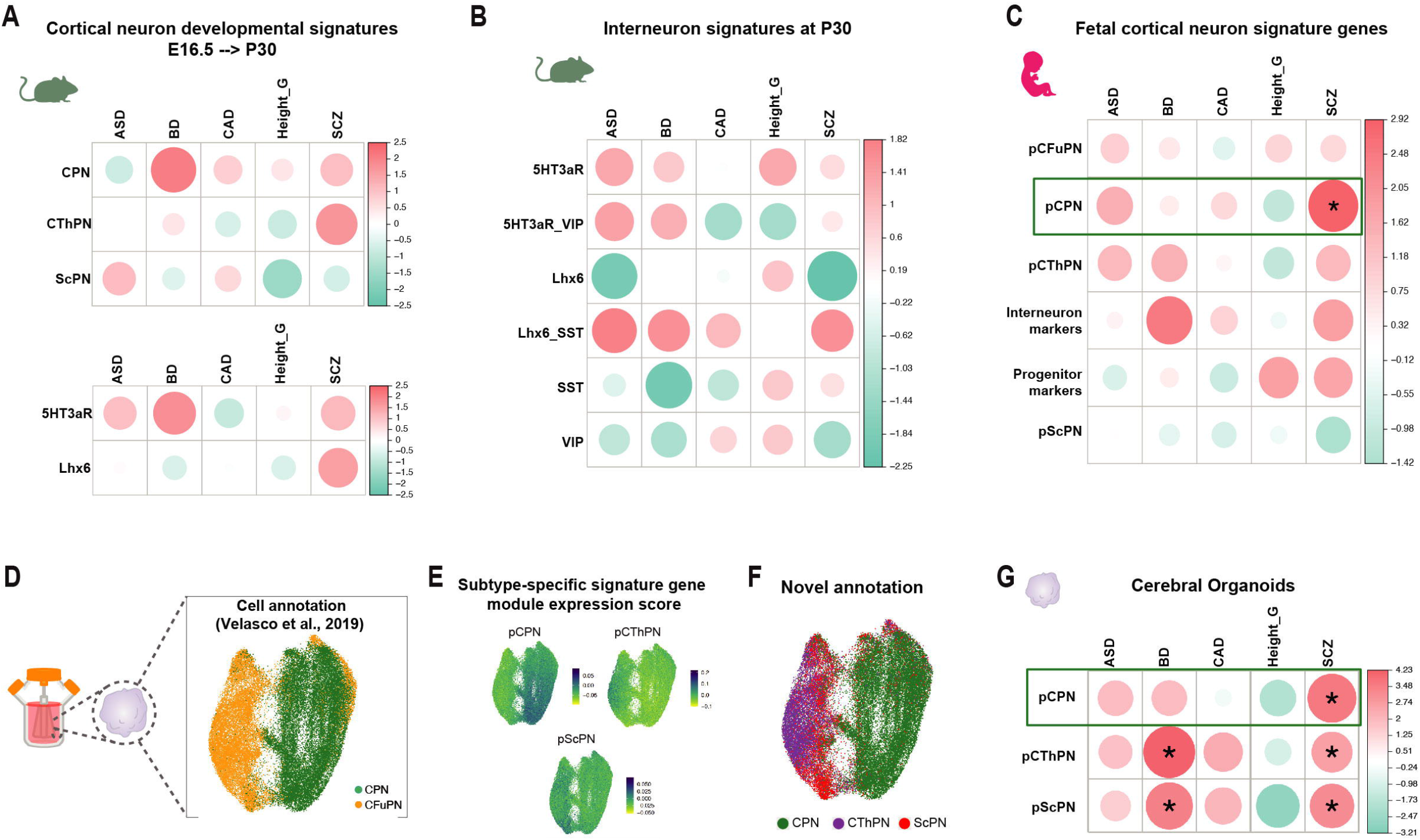
Linkage Disequilibrium Regression score for GWAS hits of neurodevelopmental and neuropsychiatric disorders in human and mouse cortical neuron signatures. Genetic correlations (estimated by LD Score Regression) between mouse (**A, B**) and human (**C**) neuronal subtype-specific signature genes and Height, Coronary Artery Disease (CAD), Autism Spectrum Disorder (ASD), Schizophrenia (SCZ), Bipolar Disorder (BD). ^*^represents significant genetic correlation. (**D-F**) Computationally extracted PN subtypes form the 3 month organoids dataset previously published^52^ were re-annotated using the PLIER algorithm which employed our signature gene sets (**E-F**). (**G**) Genetic correlations (estimated by LD Score Regression) between 5% top expressing genes of cortical organoid subtypes at 3 mo in vitro and Height, Coronary Artery Disease (CAD), Autism Spectrum Disorder (ASD), Schizophrenia (SCZ), Bipolar Disorder (BD). * represents significant genetic correlation. 5% top expressed genes and sum statistics of disease traits are listed in Table S10. Abbreviations: pCPN, putative Callosal Projection Neurons; pScPN, putative Subcerebral Projection Neurons; pCThPN, putative CorticoThalamic Projection Neurons; pCFuPNs, putative Corticofugal Projection Neurons; Im CPN, immature CPN.

By contrast, when we analyzed signature gene sets from the human samples, a significant and specific enrichment for SCZ was detected in one neuron class, CPNs (Figure 5C and Figure S10). These data indicate that developmental signatures of human, but not mouse, CPNs are enriched for genes associated with genetic risk for schizophrenia, unearthing a previously unappreciated species-specific association of this neuropsychiatric disorder with a defined cortical neuronal subtype.

We next tested whether this disease associations held true in a model of human corticogenesis, human cortical organoids^36,49-51^. Because manual annotation using small lists of known marker genes had not previously been able to discriminate between corticofugal subtypes in organoids^49,52^, we first applied our gene signatures from human projection neuron subtypes (CPNs, CFuPNs, CThPNs, and ScPNs) to infer neuronal subtype identity in our previously published single-cell RNAseq dataset of human cortical organoids^52^.

We used our human developmental signatures as input to Pathway-Level Information Extractor (PLIER)^53^, a matrix decomposition technique that optimizes latent variables using prior biological knowledge to assign each cell a new identity (see Methods). Among the excitatory neurons, we were able to clearly resolve CThPNs and ScPNs in the organoid dataset, as well as the previously-annotated CPNs (Figure 5D-F, Figure S10A-B). We then performed differential gene expression analysis to define the molecular subtype-specific signatures of organoid-derived PN subtypes (Table S10). We validated that these newly annotated ScPNs and CThPNs were enriched for known subtype-specific signature genes (i.e. LMO3, TLE4, PPP1R1B for CThPNs, LDB2, CRYM, ETV1 for ScPNs).

We next investigated whether CPNs produced within human cortical organoids recapitulated the same disease association with SCZ found in endogenous human fetal neurons. We included the DEG sets emerging from our PLIER analysis of organoid PN subtypes (Table S10), or the top 5% of DE genes (by signed t-score ranking) for each subtype, and partitioned heritability analyses to the associated genomic loci^48^. We found enrichment for SCZ risk genes in organoid CPNs, consistent with our findings on human fetal CPNs. Of note, organoids also revealed association of SCZ with organoid ScPNs and CThPNs. We also found significant enrichment for BD risk genes in organoid ScPNs and CThPNs, and neuroticism risk genes in organoid CPNs and CThPNs (Figure 5G and Figure S11), possibly reflecting a different maturation stage of the organoid neurons compared to the endogenous ages we tested.

These data identify a species-specific susceptibility of a defined neuronal subtype, callosal projection neurons, for schizophrenia. This association is not observed in mouse neurons and is identifiable during midgestation of the human fetal cerebral cortex, pointing to an early developmental effect that precedes the earliest clinical manifestations of the disease, during late human adolescence^54^.

## Discussion

The emerging wealth of genetic information on neurodevelopmental and neuropsychiatric disorders offers an opportunity to link disease-risk associated genes to the specific cell types and developmental processes they may affect. Due to the diversity of cell types present in the brain and the dynamic nature of their gene expression programs over time, attempting such associations requires a detailed understanding of the transcriptional profiles of different cell classes over multiple developmental stages.

Here we present a high-resolution, longitudinal transcriptional atlas encompassing multiple neuronal subtypes of the neocortex in both mouse and human. The data unearths notable features of the molecular programs associated with acquisition of cell type identity and neuronal diversification in the neocortex. For example, it is evident that gene signatures defining neuronal classes vary across time, indicating that the molecular identity of a neuron cannot be defined by its molecular makeup at any one age; rather, each type is represented by a collection of transitional states unfolding as development progresses.

Cross-species comparison also highlighted salient features of human brain development. Our finding that mid-gestation human fetal neurons are most similar to P1 mouse subtypes, points at accelerated cortical neuron development relative to organismal development in humans. In addition, human PN classes shared only 10-20% of their genes with their mouse counterparts, with long non-coding RNAs and unannotated loci being highly enriched in the human-specific signatures. Beyond supporting a role for non-coding loci in human brain evolution^55, 56^, the data are intriguing in the context of human disease. Genomic studies have identified the majority of risk variants associated with SCZ within non-coding regions of the genome, suggesting that they alter risk by changing levels of gene expression or splicing^57^.

Prior studies on adult cerebral cortex have suggested a common, pan-neuronal and glutamatergic enrichment for SCZ risk variants^1-11,48,58,59^. Our association data, across multiple types of neurons in mouse and human neocortex, now show that there is both neuron-type and species-specific enrichment for SCZ-associated variants in signature “core” genes of a single PN subtype of the human fetal cortex: callosal projection neurons. Structural changes in the corpus callosum and reduced spine density in layer II/III neurons have been reported in schizophrenia patients and experimental models^60-62^, consistent with this new association. This is also interesting in light of our finding that human CPNs retained the least conservation of molecular signatures compared to mouse, showing the largest enrichment of human-specific molecular features among the neurons sampled.

Collectively, the data point to developing human callosal projection neurons as a potential target cell for functional investigation of genetic risk factors in SCZ pathology, and contributes to decode the complex cellular and molecular underpinnings of schizophrenia.

## Acknowledgments

We thank Josh Huang for providing the T*cerg1l-2A-CreER* mouse line and Hannah Monyer for providing the *5HT3aR-GFP* mouse line. We are grateful to Davide Cacchiarelli for his advice on analysis of transcriptional data, Zachary Trayes-Gibson and Nathan Curry for their outstanding technical support, and Jennifer Couget for help in library preparation. We thank Stefano Stifani for providing the anti-TLE4 antibody, and all members of the Arlotta and Lodato lab for insightful comments and suggestions.

## Author contributions

E.Z., S.L. and P.A. conceived all experiments. E.Z. and S.L. performed all experiments. V.M., K.K., S.M., L.J.B., A.B., B.N., J.Z.L., and M.J.Z., performed data analysis. H.C., P.O., and C.G. contributed to sample purification and preparation. J.R.B. assisted with preparation of graphs, Figure s and with he writing of the manuscript. The work was supervised by P.A., S.L., M.J.Z., and J.Z.L. The manuscript was written by E.Z., S.L., and P.A. with contribution from all authors.

## Data and materials availability

The data for this project has been deposited in GEO under accession [<u>TBA</u>].

## Methods

### Animals

All animals were handled according to protocols approved by the Institutional Animal Care and Use Committee (IACUC) of Harvard University. All mice were maintained in standard housing conditions on a 12-h light/dark cycle with food and water ad libitum. No more than four adult animals were housed per cage. Both females and males were included in the study. Time pregnant CD1 females have been purchased at Charles River Laboratories and embryonic and postnatal litters have been collected at the desired time points (E16, E18, P1, P3, P7, P30). All the transgenic mouse lines used in this study were imported and housed in our Animal Facility: Vip-IRES-cre (Jax Stock No 010908); Sst-IRES-cre (Jax Stock No 018973); Lhx6-GFP (Lhx6-EGFP)BP221Gsat)^63^; 5HT3aR-GFP^64^ kindly provided by Hanna Monyer; and T*cerg1l-2A-CreER*^*34*^, kindly provided by Joshua Huang, was crossed with nuclear reporter line SUN1-sfGFP-myc-pA (Jax Stock. 021039).

Sequences of PCR primers employed to genotype reporter mouse lines are as follows: for 5HT3aR-GFP we used a common Forward (FW) primer GCAAGATGTGACCAAGCCACCTATTT and as Reverse (RV) TGAACTTGTGGCCGTTTACGTCG for the mutant and CAGCCCTCAGCCCTTTGAGACTTAAG to detect the wt; for Lhx6-GFP: FW GCTGAAGCACTGCACGCCGTAGG and RV GTTTGTCGGGACCTTCTTCA; For SST-Tom: TCTGAAAGACTTGCGTTTGG FW for the wt, TGGTTTGTCCAAACTCATCAA FW to detect the transgenic, and GGGCCAGGAGTTAAGGAAGA as a common RV primer; For VIP-IRES-Cre and T*cerg1l-2A-CreER* GTCCAATTTACTGACCGTACACC and GTTATTCGGATCATCAGCTACACC.

### Tissue samples

#### Mouse samples

For each biological replicate of the bulk sequencing experiment, the somatosensory cortex was dissected from one litter of mouse embryos or pups (six to ten pups per litter) or from 8-10 littermates P30 mice. For single-cell analysis, we pooled cortical regions (motor, somatosensory, auditory, visual) at distinct anterior-posterior locations (A-P). 5-6 Tcerg1l-2A-CreER P56 mice, previously injected intraperitoneally with a single dose of Tamoxifen (100□μl at 5□mg□ml^−1^) at P7. The Allen Mouse Brain framework version 3 ontology^65^ was used to define cortical primary areas.

#### Human fetal samples

De-identified human fetal tissue samples (GW16-GW21) were obtained from ABR upon patient consent in strict observance of the legal and institutional ethical regulations. Protocols were approved by the Harvard Institutional Review Board. Fetal cortices were microdissected on ice in HibernateA and dissociated into single cell suspension as previously described^66^.

### Tissue processing for cell and nuclei sorting

Enzymatic (Papain; Worthington Biochemical Corporation) and mechanical digestion were used to dissociate cortex into single cell suspension, following manufacturer’s protocol. Briefly, tissue samples were cut into small pieces and placed in a vial containing a pre-warmed solution of Papain and triturated following manufacturer protocol. After removing the dissociation media, cells were resuspended in PBS and processed for intracellular staining and FACS-sorting^18^. Somatosensory corteces obtained from P30 mice were dissociated into single cell suspension by enzymatic and mechanical digestion using the Papain dissociation kit, according to the manufacturer’s protocol (Worthington, cat. #LK003153). For nuclei preparation, somatosensory, motor, auditory and visual cortex from wild-type animals at P56 was dissected and dissociated. Each library was made from tissue pooled from at least 8 animals, and a balanced sex ratio was used. Tissue dissociation was performed as previously described, and live cells were isolated by FACS sorting as DAPI-negative, Vybrant DyeCycle Ruby (Thermo Fisher)-positive events. Libraries were prepared using the 10x Genomics Chromium Single Cell 3’ kit v2 (10x Genomics) according to the manufacturer’s protocol.

### FACS-purification of mouse and human cortical projection neurons

The protocol for intracellular staining and RNA isolation was previously described^18^. Single cell suspension was centrifuged for 5 minutes at 250g. Cell pellet was resuspended in 4% paraformaldehyde (Electron Microscopy Science), 0.1% saponin, and 1:25 RNasin in PBS (2×10^7 cells/ml) and incubated at 4°C for 30 minutes. Cells were pelleted at 3000g for 30 minutes, washed twice with wash buffer (0.1% saponin, 0.2% BSA, 1:100 RNasin in PBS), resuspended in primary antibody (0.1% saponin, 1% BSA, 1:20 RNasin in PBS), and incubated for 30 minutes at 4°C. Cells were then washed twice, incubated in secondary antibody for 30 minutes, washed twice more, and resuspended in PBS with 0.5% BSA and 1:40 RNasin for FACS purification on a BD FACS Aria II into PBS with 0.5% BSA and 1:40 RNasin. RNase free BSA was from Gemini Bio-Products. Primary antibodies were mouse anti-□SATB2 1:250 (Abcam), rat anti-CTIP2 1:500 (Abcam), and rabbit anti TLE4 1:2000 (gift from S. Stifani)^30^. Secondary antibodies were goat anti-mouse A488, goat anti-rat A546, and goat anti-rabbit A647 at 1:1000 (Molecular Probes). Appropriate gates for FACS were set based on relative levels of SATB2, CTIP2, and TLE4 expression to isolate CPN, ScPN, and CThPN, as previously described^18^.

### FACS-isolation of mouse interneuron classes

Single cell suspensions obtained from somatosensory cortices isolated from interneuron reporter lines (Vip-IRES-cre; Sst-IRES-cre; Lhx6-GFP; 5HT3aR-GFP) were analyzed by FACS and interneuron classes were FACS-purified based on the expression of the reporter gene. Appropriate gates on FACS were set based on wild type cortex, which do not express the reporter gene.

### FACS-isolation of mouse cortical nuclei

Single nuclei suspensions obtained from primary cortices isolated from *Tcerg1l-*2A-CreER P56 mice were analyzed by FACS and Tomato positive neurons isolated. Appropriate gates on FACS were set based on wild type cortex, which do not express the reporter gene.

### Immunohistochemistry

Mice were anesthetized and transcardially perfused with ice-cold PBS followed by ice-cold 4% paraformaldehyde in PBS. Dissected brains were post-fixed overnight in 4% paraformaldehyde at 4 °C, and stored in PBS-Azide 0.0025%. Brains were processed as previously described^23,30^. Vibratome brain slices were incubated with blocking media (8% goat serum, 3% BSA, 0.3% Triton in PBS) for 1hr, then with primary antibodies overnight at 4°C. Secondary antibodies were applied with 1:800 dilution in the blocking media and incubated for 2hr at room temperature, after washes. DAPI staining was performed for 3 mins before mounting with Fluoromount G (Invitrogen, #00-4958-02). The antibodies: Chicken anti-GFP antibody (ab16901, 1:500; Millipore), Mouse anti-Satb2 (ab51502, 1:50; Abcam), Rat anti-Ctip2 (ab18465, 1:100, Abcam), Rabbit anti-Tle4 (a kind gift from Stefano Stifani). Appropriate secondary antibodies were from the Molecular Probes Alexa Series. All images were acquired using a Nikon Eclips 90i fluorescence microscope and analysed with Volocity v6.0.1 software. Confocal images were obtained using an LSM 700 inverted confocal microscope (Zeiss) and analysed with a Zen Blue/Black 2012 Imgae-processing software.

### RNA extraction, library preparation, and sequencing

RNA was extracted from molecularly identified PN subtypes using the RecoverAll Total Nuclear Isolation Kit (Thermo Fisher Scientific) following manufacturer’s instruction except for crosslinking reversion, which was performed by incubating the nuclear pellet in Digestion Buffer and Protease mixture (100 μl buffer and 4 μl protease) for 3 h at 50°C. Isolated interneuron subtypes were subjected to RNA extraction with Trizol, following manufacturer’s protocol. RNA was quantified using Nanodrop and Qubit and quality assessed by Agilent 2100 Bioanalyzer. Purified RNA served as input for the cDNA library preparation with SMART-Seq v.4 Ultra Low Input RNA kit (Takara Bio, Kusatsu, Japan) and Nextera XT DNA Library Prep Kit (Illumina, San Diego, CA, USA) according to the manufacturer’s protocol. The cDNA library fragment size was determined by the BioAnalyzer 2100 HS DNA Assay (Agilent, Santa Clara, CA, USA). The libraries were sequenced as paired-end reads on HiSeq 2500 or NextSeq 500 to retrieve 1.02 ×10^8^ mapped reads. For single nuclei RNA sequencing, libraries were prepared using the Chromium Single Cell 3’ Reagent Kit v2 (10x Genomics). Briefly, FACS-purified nuclei were loaded into a channel of the Chromium single cell 3’ Chip, following manufacturer’s protocol, and partitioned into droplets in the Chromium Controller, before library construction. Libraries were then deep-sequenced to approximately 50,000 reads/nucleus.

### Computational analysis

Computational analysis was performed using R package version 3. More specifically, we used v. 3.4.4 for the analysis of bulk sequencing, v. 3.6.1 for the analysis of single cell sequencing, and v3.6.0 for the analysis of single nuclei sequencing.

### Quality control and data processing for bulk sequencing

RNA-Seq reds were mapped using STAR/2.5.0^67^with the following settings:

--outFilterType BySJout --outFilterMultimapNmax 20 --outFilterMismatchNmax 999 --outFilterMismatchNoverReadLmax 0.04 --alignIntronMin 20 --alignIntronMax 1000000 --alignMatesGapMax 1000000 --alignSJoverhangMin 8 -- alignSJDBoverhangMin 1 --sjdbScore 1 --quantMode GeneCounts

Against the GRCh38 or mm10 genome respectively using Gencode based gene annotation version 24 for human and 16M for mouse. Raw gene level quantification from STAR was used for all further analyses and Input to DESeq. Technical replicates were merged and afterwards conditional quantile normalization from cqn R package version 1.24.0 was applied to all three data sets, correcting for effects of sequencing depth, gene length and GC content. GC content information was obtained from ENSEMBL, using homo sapiens data set version 83 and mus musculus data set version 91. Gene length was defined as the average length of all listed transcripts for each gene, based on the gencode ENSEMBL transcript annotations version 24 for human and version M16 for mouse. Subsequently, batch effects introduced by flow cell association were corrected for with ComBat from sva package version 3.26.0 in both projection data sets. ComBat was not applied to mouse interneurons because sequencing flow cell and conditions of interest were confounded for this data set. In order to validate sample identity and quality, dimensional reduction and further analyses were implemented. Principal component analysis was conducted based on the prcomp function from stats package version 3.4.4 and used to check the first components explaining the majority of the observed variance for outlier samples. To further support the sample selection process, classical multidimensional scaling based on stats cmdscale function was used. Results for two different distance metrics were evaluated, euclidean distance and centered Pearson. Additional sample identity confirmation was attained through clustering of the dissimilarity matrices based on hclust and ward.D2 criterion for agglomeration. Lastly, samples were validated by checking the expression of a previously established set of known subtype marker genes. The final curated data sets encompassed 48 samples for human projections, 43 for mouse projections and 31 for mouse interneurons.

### Gene signature set definition

Differential expression was tested for with the DESeq2 package v. 1.18.1. Normalization factors were derived from conditional quantile normalization and added to the DESeq2 object, correcting for transcript length, sequencing depth and gc content. For mouse and human projection neuron data sets, flowcell identity was considered as a covariate in the design formula. The batch distribution of the mouse interneuron samples did not allow for inclusion of flow cell information in the design formula for this data set. The significance threshold for adjusted p values was set to 0.01. Subsequently, the resulting list of differentially expressed genes was filtered, using an empirical approach to identify meaningful parameters to pinpoint cell type specific marker genes. To that end, reference lists of known marker genes for projection and interneurons were used to define thresholds on normalized gene expression, filtering based on average log2 fold change and cell type specificity (defined as entropy). Thresholds were chosen so that the marker genes present in our preliminary lists were retained in the final sets as well. The results are listed in Table 2. Resulting signature genes were then clustred using PAM (pam() function from R package cluster v. 2.1.0). The input was defined as the average gene expression per subtype and age for each gene retained after filtering. Optimal k were determined using gap statistics as implemented in clusgap version 2.1.0. Genes resulting from human (k 23), mouse projection (k 7) and the two different lists of mouse interneuron genes (k 18 for 5HT3aR vs Lhx6 and k 24 for all four subtypes at P30) were all clustered separately. To each of the resulting clusters a subtype identity was assigned. Different clusters assigned to the same subtype showed different temporal expression patterns.

Two strategies were implemented to merge the clusters to obtain the final gene sets. First, all clusters of the same subtype were merged together to obtain “subtype sets”. Second, all clusters within a subtype were clustered hierarchically (centered Pearson) using the overall average expression of all genes within a cluster for each subtype and age. All clusters derived from the same data set were merged according to the same distance cutoff. This second strategy was not applied to the mouse projection clusters, because they were already bigger in size than the clusters derived from the other two data sets where the chosen k were generally larger. Instead, all 7 clusters were treated as individual gene sets in the disease association analysis. All mouse derived sets were mapped to one2one human orthologues. The results are listed in Table 3.

### Single-nuclei RNA sequence analysis

Single-nuclei RNA-seq reads were aligned to mm10 pre-mRNA reference, and gene expression matrix was obtained using CellRanger software v3.1.0 with the default parameters. R v3.6.0 and Seurat package v3.1.2 was used to perform downstream analyses. During the analysis, cells from the piriform region were removed, in order to focus our study on the ScPNs. Low-quality cells (with percent mitochondrial gene expression (percent.mt) >0.5%) were removed from the analyses. SCTransform was performed to normalize and scale the gene expression matrix. During the SCTransform, the number of UMIs per cell and percent.mt were treated as variables to regress out. Since we want to remove sex effects linked to the presence of chromosomes, X and Y chromosome genes were removed from the list of variable genes identified. The resulting list of variable genes was used to perform the principal component analysis. Using the top 30 principal components (ordered by the fraction of the total variance explained) and Louvain clustering algorithm (FindClusters function in Seurat with resolution set to 0.7), 16 different cell clusters were identified. This dimension of 30 PCs is further reduced to two Uniform Manifold Approximation and Projection (UMAP) dimensions (RunUMAP function in Seurat with dims parameter set to top 30 PCs) for visualizations. Using known marker genes (listed in Table: Markers), clusters were classified into four broad cell types: ScPN, upper layer CPN, inhibitory neurons and low quality.

#### Markers

| ScPN | UL CPN | Inhibitory Neurons |
| --- | --- | --- |
| Fezf2 | Rorb | Synpr |
| Parm1 | Cux1 | Sst |
| Tcerg1l | Cux2 | Gad1 |
| Bcl11b |  | Gad2 |

### Finding area-specific markers of ScPN

ScPNs were selected from the broad cell type annotation, and were processed separately by applying the same workflow described above with percent.mt cutoff of 0.2%. Treating brain slice/functional area annotations as cell clustering, differentially enriched genes for each section were identified using MAST v1.10.0^68^ (using FindAllMarkers function with test.use parameter set to MAST, latent variable set to the number of UMIs and percent.mt, min.pct set to 0.3). To group DEGs together by their patterns of expression, we first computed the average expression of these genes in each brain slice/functional area (using the AverageExpression function in Seurat). Z-scores were then computed to find enriched or depleted genes in each region. To group differentially expressed genes into groups based on their expression pattern, K-means clustering (Hartigan-Wong algorithm implemented in R base package named stats) was used. The number of clusters was determined by benchmarking several peaks from the gap statistics result (obtained from clusGap function in cluster package v2.0.8 with parameters K.max=20 and B=60).

### Organoid cell type re-annotation

We took the dataset produced in Velasco et al., 2019^52^ to test the performance of our gene signatures to accurately identify subtypes of PNs. To do so, we selected cells labeled as CPNs and CFuPNs according to the published annotation and processed only those cells. Normalized count matrix containing the selected cells was taken from the published dataset and processed using Seurat package in R using the default parameters unless otherwise specified. Variable genes were found using FindVariableFeatures function, and the ScaleData function was used to scale the dataset. Principal component analysis was performed using RunPCA function on the scaled data, and the top 18 PCs were chosen based on an elbow plot generated using ElbowPlot function. RunUMAP function was used on the top PCs to visualize the variation in cells. As the original authors did, Harmony (version 1.0) was used to remove the batch effect among different cell lines and replicates. The dataset contained HUES66 and PGP1 cell lines, and PGP1 line had two replicates. To identify specific PN subtypes (CPN, CThPN and SCPN) in this processed organoid dataset using our signatures gene lists, we applied PLIER (PLIER function from PLIER package version 0.99.0) specifying our signature lists as the prior knowledge (gene sets). Among the resulting latent variables that align to the prior knowledge, only the ones with AUC > 0.5 and p value < 0.005 were considered in the reannotation. The new annotation of each cell was chosen by the latent variable with the highest cell loading.

### Validation of human cell type signature genes in a published dataset

We validated our curated human gene modules with a previously published dataset Nowakowsky et al., 2017^43^. This scRNA-seq dataset was processed from the downloaded gene count matrix, using a similar downstream pipeline as specified in the single-nuclei RNA sequence analysis. Note that percent.mt was not used to filter out the cells and when scaling the dataset, because the original authors had already removed mitochondrial genes by only keeping genes expressed in at least 30 cells. Using our gene signature modules curated for each neuronal subtype of interest (CPN, CfuPN, CThPN and ScPN), we calculated gene signature scores to identify clusters expressing these modules at high levels. AddModuleScore() function from Seurat was used to compute gene signature scores, setting the control parameter to 30. Based on our cell cluster definition and the module scores, we applied cluster-based re-annotation strategy, where cell type annotations were reassigned if cells from a specific cluster showed enrichment with one of our modules. Enriched clusters were determined by looking at both the overall module scores in clusters and the expression of individual genes.

### Ortholog identification

Human orthologs for mouse genes were obtained based on the mus musculus gene data set from ENSEMBL version 91 and biomaRt package version 2.34.2. Genes of mouse homology types (one to many; many to many) mapping were excluded and only “one-to-one” orthologs were considered for downstream analyses. Mitochondrial genes were excluded resulting in a total set of orthologs encompassing 15,974 genes. 6,336 of those genes were found to be differentially expressed both in human and in mouse projection neurons. 1,804 orthologs passed gene list filtering in at least one species and were thus identified as potential signatures and 302 of those passed in both species, with 234 of them being included in the final signature sets. 165 of those were assigned to the same neuronal subtype signature in both species and consequently identified as convergent signatures.

### Interspecies comparison analysis

Raw counts for 48 human and 43 mouse samples were combined by retaining expression values for 15,974 ENSEMBL human mouse orthologs. Based on the combined count data, a DESeqDataSet object was created. Neuronal subtype and age were considered in the design formula as a joint variable in addition to the sequencing flow cells. Conditional quantile normalization was applied to remove the effect of GC content, and gene length was modelled as a smooth function to account for the lengths effect on gene count. The results were set as normalization factors before estimating size factors and dispersions and fitting a negative binomial GLM with DESeq2. A blinded variance stabilizing transformation (VST) was computed for the 15,966 genes that did converge in the previous step. ComBat from sva package version 3.26.0 was run to correct for batch covariate flow cell on the basis of a parametric empirical Bayes framework. Biological replicates were summarized by their mean transformed and corrected expression values. For classical metric multidimensional scaling (MDS), analyses were implemented with R package stats version 3.4.4. Distance matrices were computed with amap version 0.8-16 choosing centered Pearson (1-corr(x,y)) as the distance metric. Supplementary Figure 9 shows MDS for all 15,966 orthologs and for the subset of 1,803 orthologs that were identified as potential signatures in at least one species. Correlation maps were created on Pearson correlated data, Figure 4 c) shows the results of correlation analysis for 302 genes that were identified as potential neuronal projection signatures both in human and in mouse.

### Functional annotation and Enrichment Map pathway analysis visualization

Pathway enrichment analysis was carried out by searching for enriched gene-sets (e.g. pathways, molecular functional categories, biological processes) in the human signature gene set versus the murine DEGs at P1 for each neuronal subtype by mapping functional information from pathways annotation collection (updated version April 2019) using gProfiler. The collection comprises Gene Ontology annotations *(Biological Process, Cellular Component, Molecular Function)* as well as Reactome, Panther Pathway, Msigdb C2 and Wikipathways terms. The resulting enrichment results were visualized with the Enrichment Map plugin for the Cytoscape network visualization and analysis software. We loaded gProfiler results using a FDR cut-off of 0.05. In these maps, each gene set is symbolized by a node in the network. Node size corresponds to the number of genes comprising the gene-set. The enrichment scores for the gene-set are represented by the node’s color intensity (green for CPN, red for ScPN, violet for CThPN and yellow for CFuPN). The color of the node upper hemisphere indicates the enrichment score for mouse gene sets, and the lower hemisphere indicates the score for human gene sets. To intuitively identify redundancies between gene sets, the nodes are connected with edges if their contents overlap by more than 50%. The thickness of the edge corresponds to the size of the overlap. The edge belonging to human dataset were represented in dark gray, those corresponding to mouse dataset were represented in light gray. We used version 1.1.0 of the Enrichment Map software in Cytoscape 3.8.0^69^.

### Gene ontology and functional enrichment analysis

From curated lists of genes, enriched biological pathways and molecular functions were identified using enrichGO function from an R package clusterProfiler (v3.14.3). We queried different databases for different organisms by setting OrgDb parameter (org.Hs.eg.db for human dataset and org.Mm.eg.db for mouse dataset, respectively) and reported only the significant terms and pathways by setting pvalueCutoff=0.05. Background list of genes were prepared for each dataset to include all the genes that are expressed in the dataset.

### LDscore regression and partition heritability analysis

Human gene coordinates corresponding to different cell types and cell type clusters (categories) were overlapped using bedtools (Version: v2.27.1-1-gb87c465) with the SNPs in the .bim file used for the computation of LD scores (1000 Genomes Phase 3 downloaded from https://data.broadinstitute.org/alkesgroup/LDSCORE/), the resulting annotation files were used to calculate the annotation specific LD scores separately for each category using LD Score Regression (LDSC) Version 1.0.0. To asses each category’s contribution to h^2, a cell type specific analysis was performed for multiple disease traits, including Schizophrenia, Autism, Bipolar Disorder, Coronary Artery Disease (Table S 10) using their corresponding GWAS summary statistics (obtained here: https://data.broadinstitute.org/alkesgroup/LDSCORE/) and a modified baseline model v1.1 which contains 52 categories. To that end, we followed the CTS workflow according the software recommendations. This modification was done to add an extra category as a control (similar to the Finucane et al. 2018^48^) that included all analyzed genes in our RNA-seq. After adjusting each category’s p-value together with the baseline model 52+1 categories to account for multiple testing, significant heritability enrichment was determined using FDR < 0.05.

