## Supplementary Information for "Human-specific enrichment of schizophrenia risk-genes in callosal neurons of the developing neocortex"

**Figure S1**

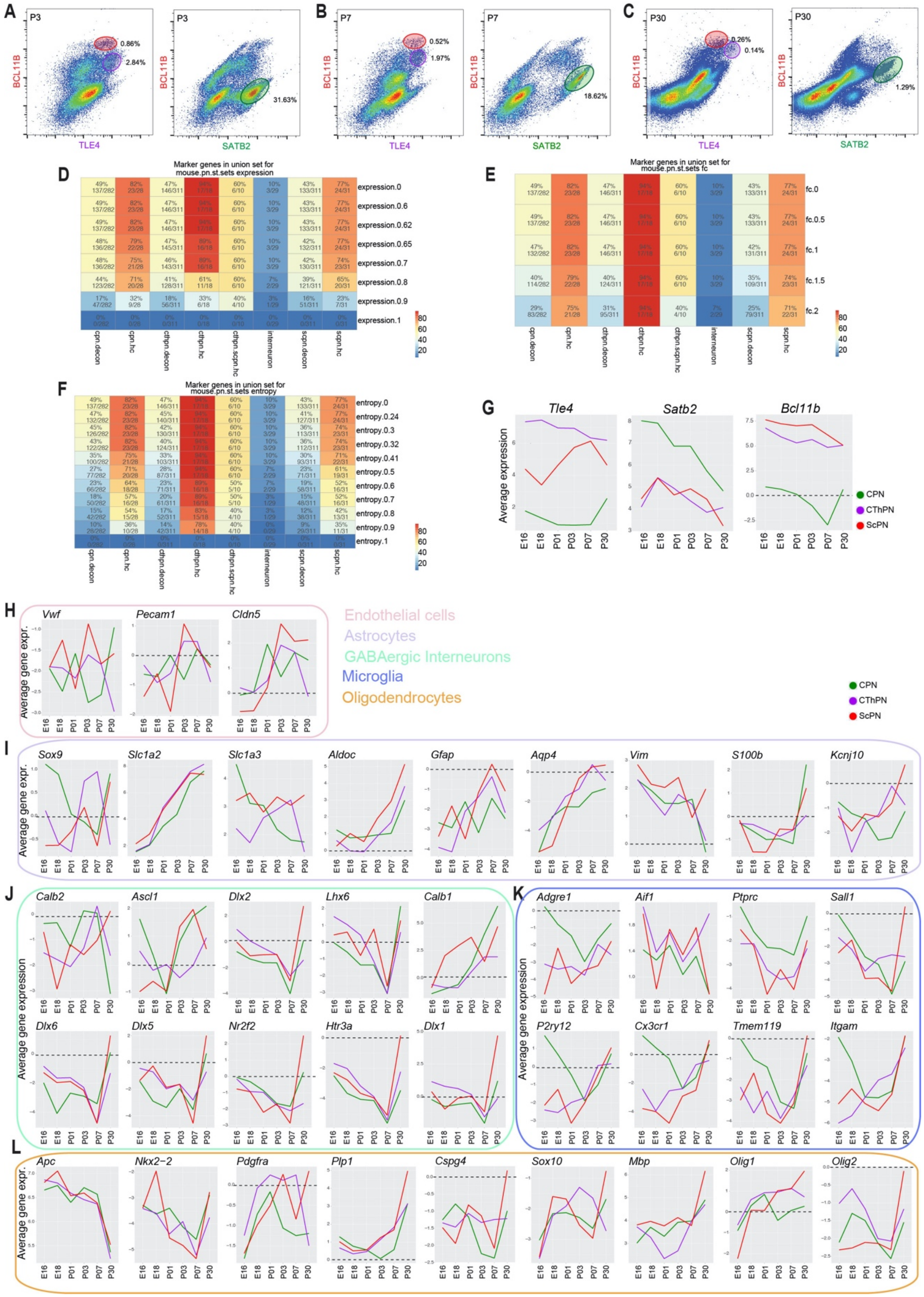
A
B
C
P3
P7
P30
BCL11B
TLE4
SATB2
0.86%
2.84%
31.63%
0.52%
1.97%
18.62%
0.26%
0.14%
1.29%
D
Marker genes in union set for mouse.pn.st.sets expression
E
Marker genes in union set for mouse.pn.st.sets fc
F
Marker genes in union set for mouse.pn.st.sets entropy
cpn.decon
cpn.hc
cthpn.decon
cthpn.hc
cthpn.scpn.hc
interneuron
scpn.decon
scpn.hc
G
Tle4
Satb2
Bcl11b
Average expression
CPN
CThPN
ScPN
H
Vwf
Pecam1
Cldn5
Average gene expr.
Endothelial cells
Astrocytes
GABAergic Interneurons
Microglia
Oligodendrocytes
I
Sox9
Slc1a2
Slc1a3
Aldoc
Gfap
Aqp4
Vim
S100b
Kcnj10
J
Calb2
Ascl1
Dlx2
Lhx6
Calb1
Dlx6
Dlx5
Nr2f2
Htr3a
Dlx1
Average gene expression
K
Adgre1
Aif1
Ptprc
Sall1
P2ry12
Cx3cr1
Tmem119
Itgam
L
Apc
Nkx2-2
Pdgfra
Plp1
Cspg4
Sox10
Mbp
Olig1
Olig2
E16
E18
P01
P03
P07
P30

**Figure S1: FACS-purification and molecular profiling of murine cortical PN subtypes**.

**A-C)** Representative FACS plots summarizing the gating strategy to identify and isolate CPNs (green, high SATB2, low BCL11B, low TLE4), ScPNs (red, low STAB2, high BCL11B, low TLE4), CThPNs (violet, low SATB2, low BCL11B, high TLE4) at (A) P3, (B) P7, and (C) P30 mouse cerebral cortices. **D-F**) Filtering parameters used to identify PN subtype-specific gene sets: (D) expression, (E) fold change, and (F) entropy. Statistical details for each figure panel are reported in Table S2. **g)** Line plots representing the expression profile of *Tle4*, *Satb2*, and *Bcl11b*, used to molecularly identify the PN subtypes. **H-L**) Line plots representing the expression profile of known markers for (H) endothelial cells, (I) astrocytes, (J) GABAergic interneurons, (K) microglia, (L) oligodendrocytes, virtually absent from purified PNs subtypes.

Abbreviations: CPN, Callosal Projection Neurons; ScPN, Subcerebral Projection Neurons; CThPN, CorticoThalamic Projection Neurons.

*Related to main Figure 1.*

### Figure S2

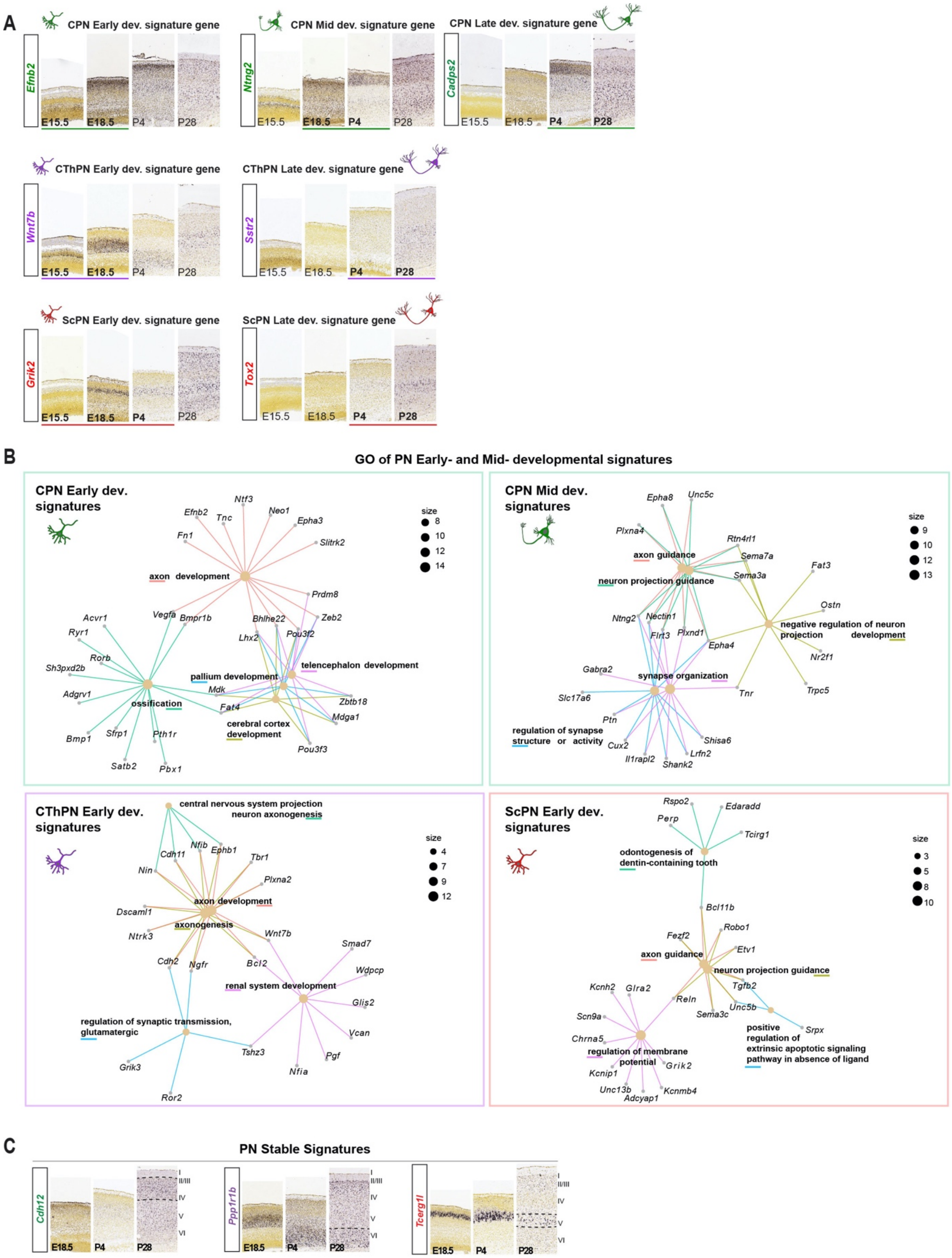
A
CPN Early dev. signature gene
Efnb2
E15.5
E18.5
P4
P28
CPN Mid dev. signature gene
Ntng2
CPN Late dev. signature gene
Cadps2
CThPN Early dev. signature gene
Wnt7b
CThPN Late dev. signature gene
Sstr2
ScPN Early dev. signature gene
Grik2
ScPN Late dev. signature gene
Tox2
B
GO of PN Early- and Mid- developmental signatures
CPN Early dev. signatures
axon development
ossification
pallium development
telencephalon development
cerebral cortex development
size
CPN Mid dev. signatures
axon guidance
neuron projection guidance
negative regulation of neuron projection development
synapse organization
regulation of synapse structure or activity
CThPN Early dev. signatures
central nervous system projection neuron axonogenesis
axon development
axonogenesis
renal system development
regulation of synaptic transmission, glutamatergic
ScPN Early dev. signatures
odontogenesis of dentin-containing tooth
axon guidance
neuron projection guidance
regulation of membrane potential
positive regulation of extrinsic apoptotic signaling pathway in absence of ligand
C
PN Stable Signatures
Cdh12
Ppp1r1b
Tcerg1l
E18.5
P4
P28
I
II/III
IV
V
VI

**Figure S2: Mouse PN subtype-specific developmental and stable signatures.**

**A)** ISH images adapted from Allen Brain Atlas of selected genes for early (*Efnb2* for CPN, *Wnt7b* for CThPNs*, Grik2* for ScPNs), mid- (*Ntng2* for CPNs), and late (*Cadps2* for CPNs*, Sstr2* for CThPNs*, Tox2* for ScPNs) subtype-specific developmental signature genes (full list reported in Table S3). **B)** GO of PN early- and mid- developmental signatures reveals enrichment of developmental processes, such as axonogenesis and forebrain development. **C)** Representative ISH images of PN subtype-specific stable signature genes: *Cdh12* for CPNs, *Ppp1r1b* for CThPNs, and *Tcerg1l* for ScPNs (modified from Allen Brain Atlas). GO terms enrichment p-values are reported in Table S4.

Abbreviations: CPN, Callosal Projection Neurons; ScPN, Subcerebral Projection Neurons; CThPN, CorticoThalamic Projection Neurons.

*Related to main Figure 1.*

### Figure S3

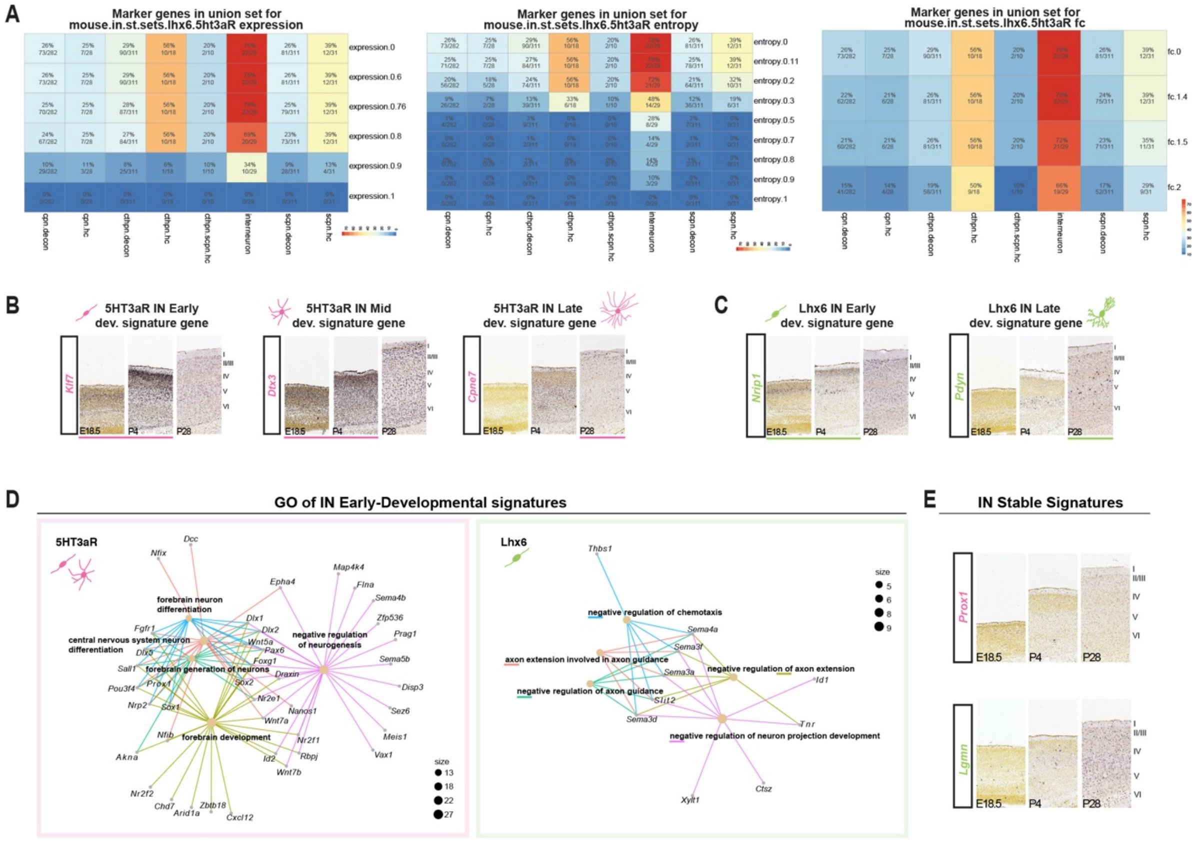


**Figure S3: FACS-purification and molecular profiling of murine cortical IN subtypes.** **A)** Filtering parameters used to identify IN subtype-specific gene sets: expression, fold change, and entropy. Statistical analysis details for each figure panel are reported in Table S2 **B-C)** ISH images adapted from Allen Brain Atlas of representative genes for early (*Ktlf*), mid- (*Dtx3*), and late (*Cpne7*) 5HT3aR- and early (*Nrip1)* and late (*Pdyn*) Lhx6-specific developmental signature genes (full list reported in Table S3). **D)** GO enriched in early- and mid- developmental signatures of IN subtypes reveals enrichment of developmental processes, such as forebrain development and neuronal differentiation. GO terms enrichment p-values are reported in Table S4. **E)** Representative ISH images of IN subtype-specific stable signature genes: *Prox1* for 5HT3aR and *Lgmn* for Lhx6 (adapted from Allen Brain Atlas).

Abbreviations: CPN, Callosal Projection Neurons; ScPN, Subcerebral Projection Neurons; CThPN, CorticoThalamic Projection Neurons; IN, interneurons.

*Related to main Figure 1.*

**Figure S4**

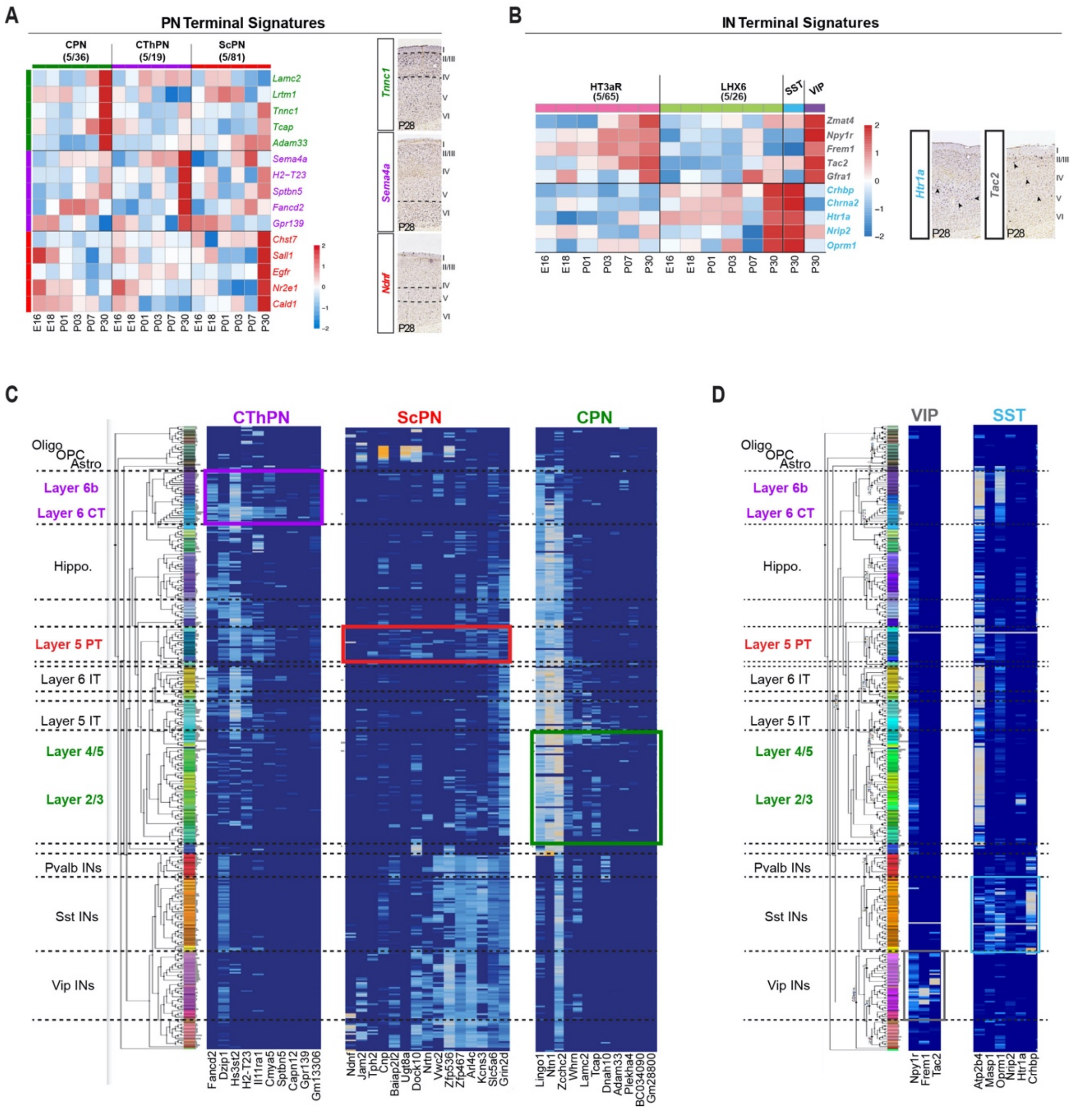


**Figure S4: Terminal signatures of murine cortical PN and IN subtypes. A,B)** Heatmap representing the expression of 5 top ranked terminal signature genes of PN (**A**) and IN. **B**) subtypes. Full list of terminal genes is reported in Supplementary Table 5. ISH images of representative terminal genes confirm their selective expression at P28 (adapted from Allen Brain Atlas) (**A-B**); **C, D**) PN (**C**) and IN (**D**) subtype-specific terminal signatures expression in published single cell data derived from P56 mouse cortex (Allen Mouse Cell Type Database[24]).

Abbreviations: E, embryonic day; P, postnatal day; CPN, Callosal Projection Neurons;

ScPN, Subcerebral Projection Neurons; CThPN, CorticoThalamic Projection Neurons; IN, interneurons; IT, Intratelencephalic; PT, Pyramidal Tract; CT, corticothalamic; Oligo, Oligodendrocytes; Astro, astrocytes; OPC, oligodendrocyte precursor cells; Hippo, hippocampus; Pval, Parvalbumin; SST, Somatostatin; VIP, Vasointestinal Peptide.
*Related to main Figure 1.*

**Figure S5**

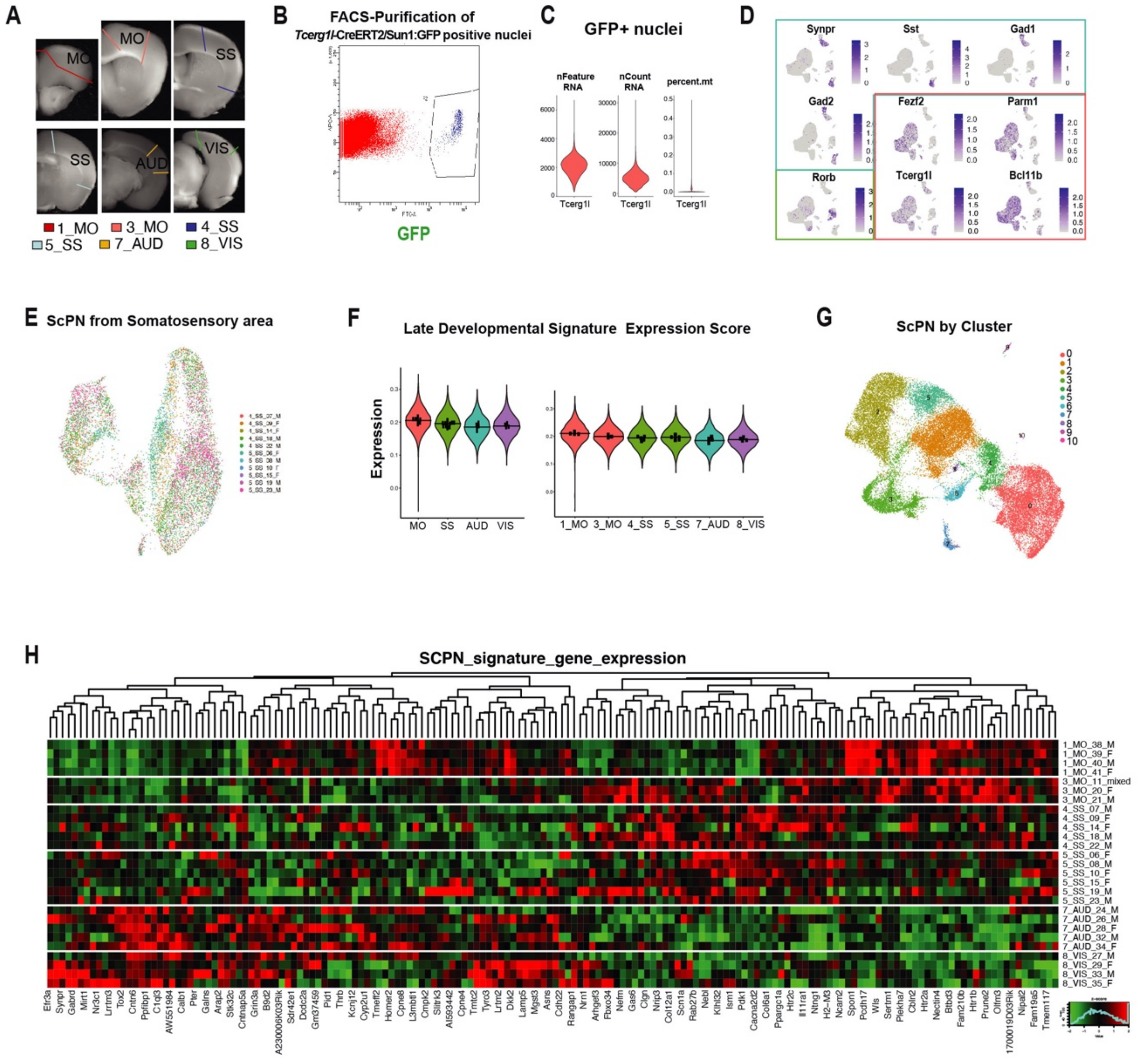


**Figure S5: Single nuclei sequencing of *Tcerg1l*-positive ScPNs from distinct mouse cortical areas and along Anterior Posterior (AP) axis at P56. A)** Sample collection strategy for layer V isolation from motor (MO), somatosensory (SS), auditory (AUD) and visual (VIS) cortical areas. Slices were collected using Allen Brain Atlas v3 "Mouse, Adult, 3D Coronal" as reference. **B**) Representative FACS plot showing gating and isolation of *Tcerg1l*-CreERT2/Sun1:GFP nuclei for downstream single nuclei RNA sequencing (snRNA seq). **C**) Quality Controls for snRNA. Metrics shown are the number of genes per nuclei (left), the number of UMIs per nuclei (center) and the percentage of mitochondrial gene expression per nuclei (right). Detailed information about samples size (number of nuclei) and QC analysis are reported in Table **S**6. **D**) Expression feature plots of selected and previously known marker genes for cortical subtypes on 45,289 GFP+ profiled nuclei. Bounding boxes are colored according to cell type identity: inhibitory neurons (light blue), ScPNs (red) and Layer IV neurons (green). **E**) UMAP plots of ScPN nuclei isolated from somatosensory cortical areas, as reported in Figure 2C. Nuclei are colored according to sample library. **F**) Violin plots showing the expression score of ScPN late developmental signatures across distinct functional cortical areas and along AP axis. **G**) Uniform Manifold Approximation and Projection (UMAP) plot of ScPN nuclei across distinct functional cortical areas and along AP axis, as reported in Figure 2D, G, showing unsupervised clustering. Nuclei are colored according to cluster. **H**) Heatmap representation of gene module expressions. Gene modules are defined by applying k-means clustering algorithm on DE genes computed by functional area in all samples. Abbreviations: P, postnatal day; CPN, Callosal Projection Neurons; ScPN, Subcerebral Projection Neurons; CThPN, CorticoThalamic Projection Neurons; IN, interneurons; Mo, Motor; SS, Somatosensory; AUS, Auditory; VIS, Visual; AP, anterior-posterior; F, female; M, male.

*Related to main Figure 2.*

**Figure S6**

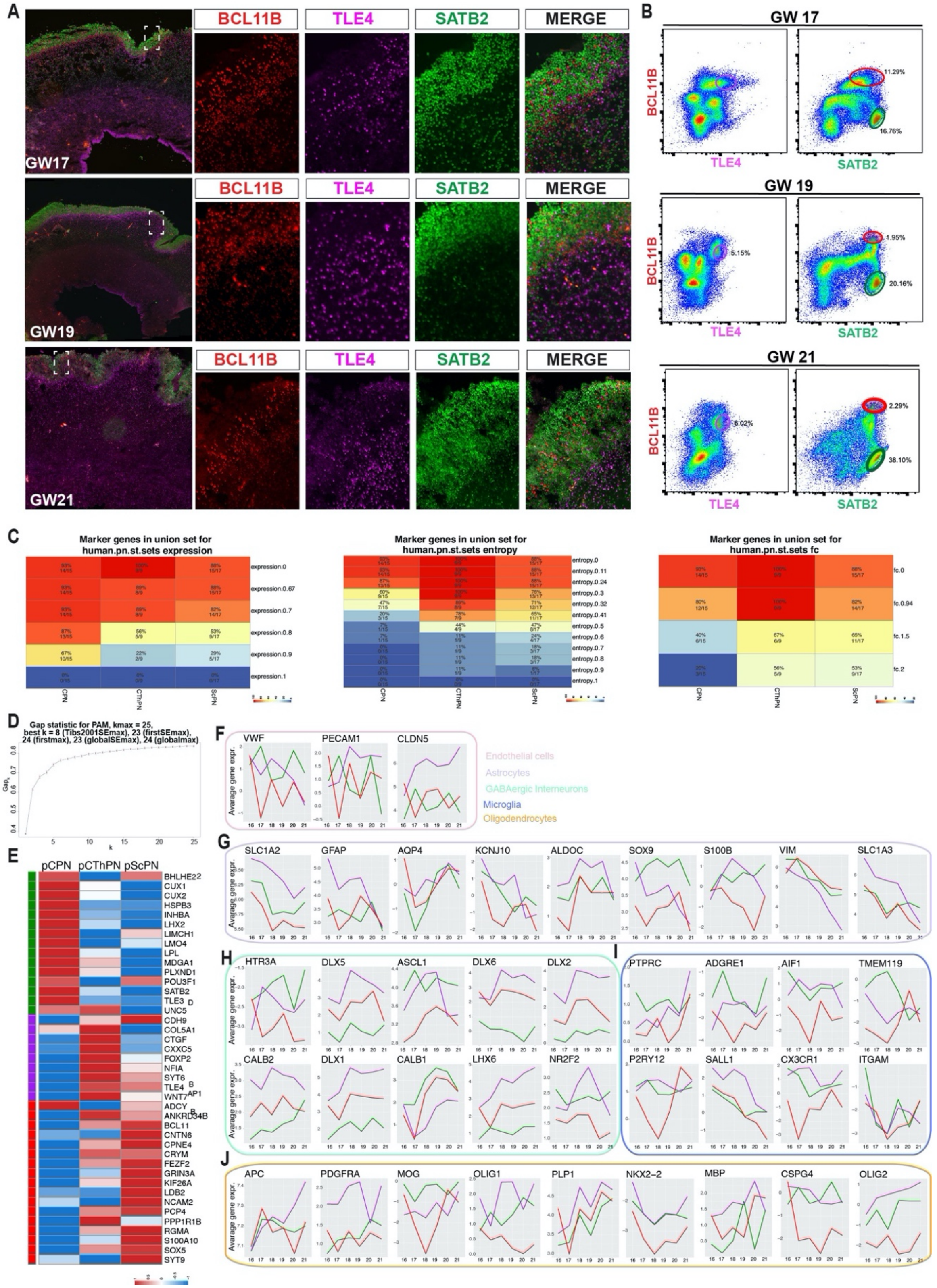

A
BCL11B
TLE4
SATB2
MERGE
GW17
GW19
GW21
B
GW 17
GW 19
GW 21
BCL11B
TLE4
SATB2
11.29%
16.76%
5.15%
1.95%
20.16%
6.02%
2.29%
38.10%
C
Marker genes in union set for human.pn.st.sets expression
Marker genes in union set for human.pn.st.sets entropy
Marker genes in union set for human.pn.st.sets fc
CPN
CThPN
ScPN
D
Gap statistic for PAM, kmax = 25, best k = 8 (Tibs2001SEmax), 23 (firstSEmax), 24 (firstmax), 23 (globalSEmax), 24 (globalmax)
E
pCPN
pCThPN
pScPN
F
VWF
PECAM1
CLDN5
Endothelial cells
Astrocytes
GABAergic Interneurons
Microglia
Oligodendrocytes
G
SLC1A2
GFAP
AQP4
KCNJ10
ALDOC
SOX9
S100B
VIM
SLC1A3
H
HTR3A
DLX5
ASCL1
DLX6
DLX2
CALB2
DLX1
CALB1
LHX6
NR2F2
I
PTPRC
ADGRE1
AIF1
TMEM119
P2RY12
SALL1
CX3CR1
ITGAM
J
APC
PDGFRA
MOG
OLIG1
PLP1
NKX2-2
MBP
CSPG4
OLIG2
Avarage gene expr.

**Figure S6**: **FACS-identification and purification of molecularly determined human PN subtypes**. **A**) Representative images of human cortices isolated from Gestational week (GW), GW17, GW19, and GW21 showing triple immunostaining for BCL11B (red), TLE4 (violet), and SATB2 (green). **B**) Representative FACS plots showing gating strategy for the isolation of distinct cortical populations at different stages, based on the expression of the three transcription factors. **C**) Filtering parameters used to identify putative PN subtype-specific gene sets: expression, entropy, and fold change. **D**) Gap statistics for *k*-clustering determination. **E**) heatmap showing the expression of known markers of distinct PN subtypes in mouse cortex. **F-J**) line plots representing the expression profile of known markers for (F) endothelial cells, (G) astrocytes, (H) GABAergic interneurons, (I) microglia, (J) oligodendrocytes.

Abbreviations: pCPN, putative Callosal Projection Neurons; pScPN, putative Subcerebral Projection Neurons; pCThPN, putative CorticoThalamic Projection Neurons.

*Related to main Figure 3.*

**Figure S7**

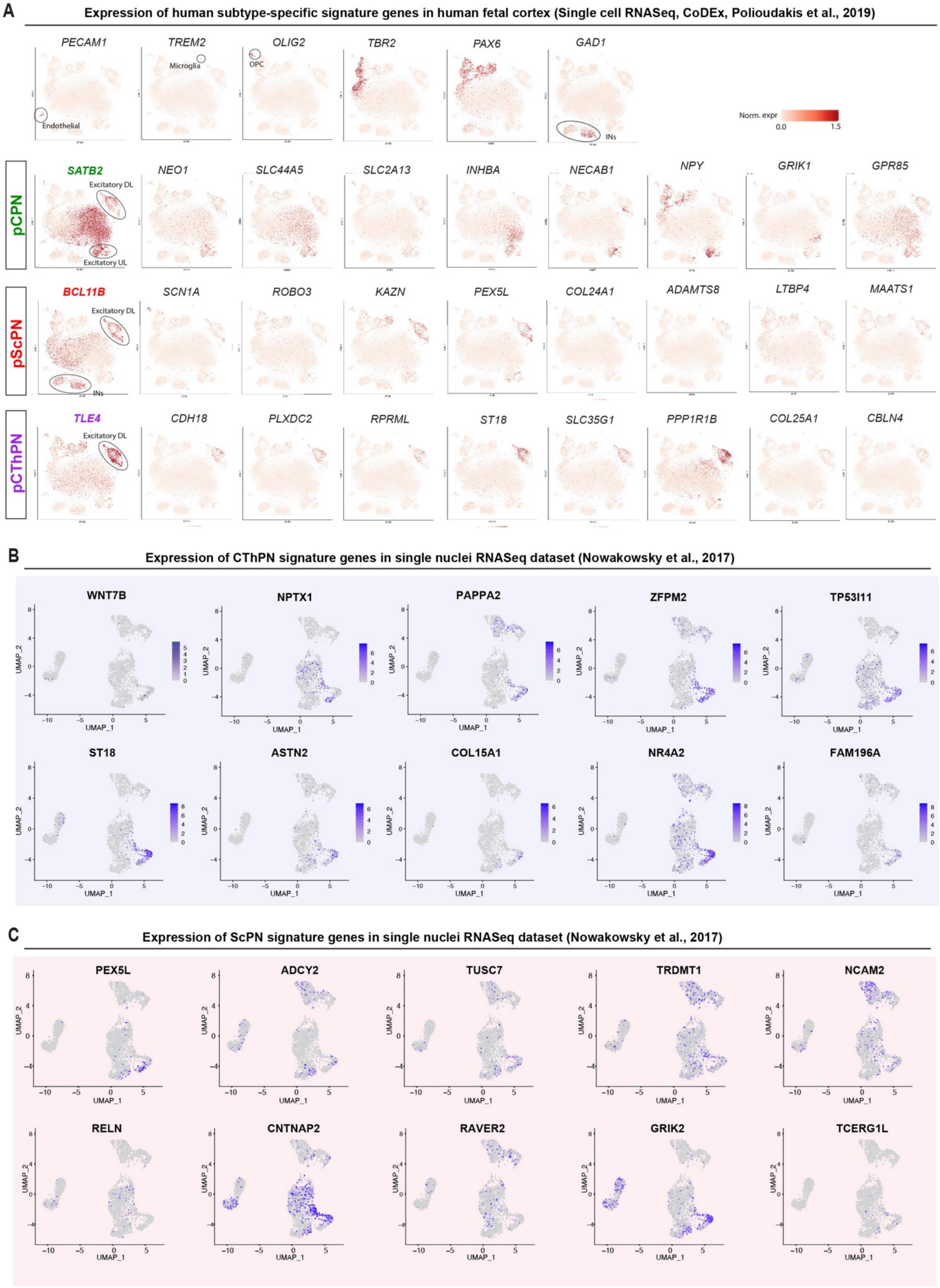

A
Expression of human subtype-specific signature genes in human fetal cortex (Single cell RNASeq, CoDEx, Polioudakis et al., 2019)
PECAM1
TREM2
OLIG2
TBR2
PAX6
GAD1
Microglia
OPC
Endothelial
INs
Norm. expr
0.0
1.5
pCPN
SATB2
Excitatory DL
Excitatory UL
NEO1
SLC44A5
SLC2A13
INHBA
NECAB1
NPY
GRIK1
GPR85
pScPN
BCL11B
Excitatory DL
INs
SCN1A
ROBO3
KAZN
PEX5L
COL24A1
ADAMTS8
LTBP4
MAATS1
pCThPN
TLE4
Excitatory DL
CDH18
PLXDC2
RPRML
ST18
SLC35G1
PPP1R1B
COL25A1
CBLN4
B
Expression of CThPN signature genes in single nuclei RNASeq dataset (Nowakowsky et al., 2017)
WNT7B
NPTX1
PAPPA2
ZFPM2
TP53I11
ST18
ASTN2
COL15A1
NR4A2
FAM196A
UMAP_1
UMAP_2
C
Expression of ScPN signature genes in single nuclei RNASeq dataset (Nowakowsky et al., 2017)
PEX5L
ADCY2
TUSC7
TRDMT1
NCAM2
RELN
CNTNAP2
RAVER2
GRIK2
TCERG1L
UMAP_1
UMAP_2

**Figure S7: Expression of Human PN subtype-specific signatures in fetal tissue at single cell resolution. A**) t-SNE plots representing the expression of human PN subtype-specific signature genes in human fetal cortex (CoDEx database, Polioudakis et al., 2019), normalized gene expression is reported on the left. **B-C**) Feature plots showing the expression of human CThPN-(**B**) and ScPN-(**C**) specific signatures in previously published single cell dataset of fetal cerebral cortex (Novakowski et al., 2017).

Abbreviations: pCPN, putative Callosal Projection Neurons; pScPN, putative Subcerebral Projection Neurons; pCThPN, putative CorticoThalamic Projection Neurons; Excitatory DL, deep layers; Excitatory UL, upper layers; OPC, oligodendrocyte precursor cells; IN, interneurons.

*Related to main Figure 3.*

**Figure S8**

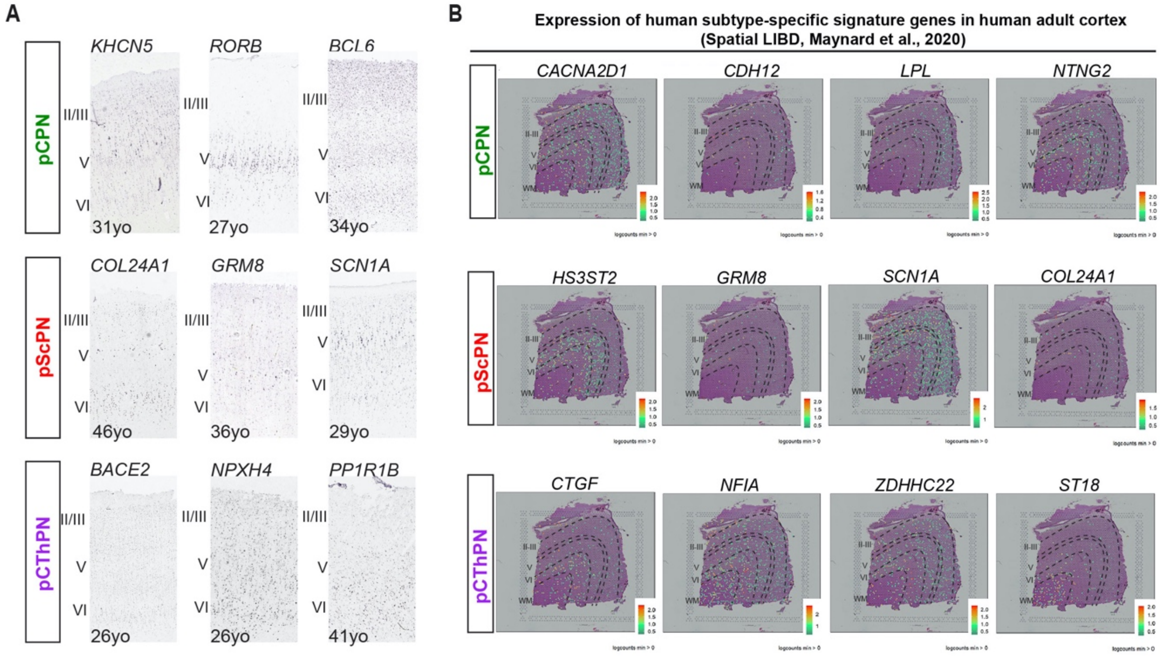


**Figure S8: Specific Expression of Human PN subtype signatures is maintained in adult cortical tissue. A**) Representative images of ISH of our PN subtype-specific genes on human adult cortex from different individuals and ages (adapted from Allen Brain Atlas). **B**) Spotplots depicting log(transformed normalized expression) (log(counts)) for sample 151673 for human PN subtype-specific signature genes on human adult cortex using Spatial Transcriptomics, LIBD database (Maynard et al., 2020).

Abbreviations: pCPN, putative Callosal Projection Neurons; pScPN, putative Subcerebral Projection Neurons; pCThPN, putative CorticoThalamic Projection Neurons; Excitatory DL, deep layers; Excitatory UL, upper layers.

*Related to main Figure 3.*

**Figure S9**

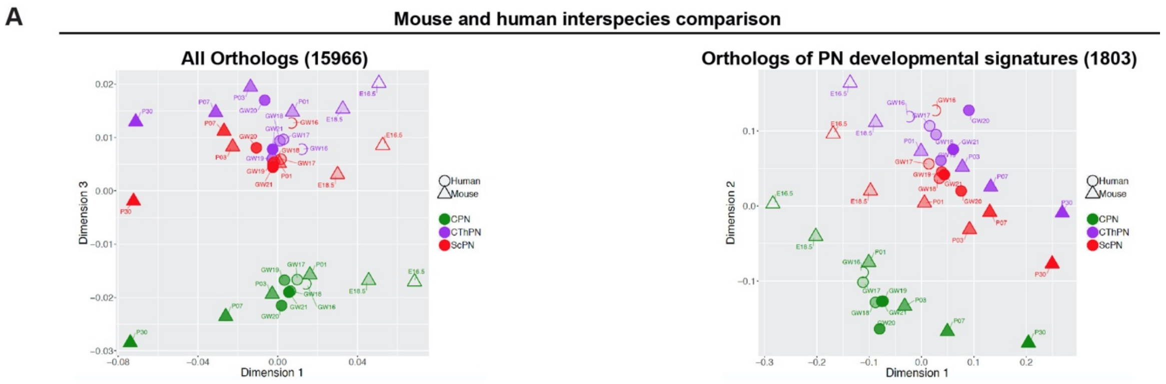


**Figure S9: Mouse and Human interspecies comparison. A**) Multiple scaling dimensionality (MDS) representing average expression of all ortholog genes in human (represented with a circle) and mouse (triangle) PN subtypes (left panel), in PN specific developmental signatures across multiple developmental stages (right panel).

Abbreviations: CPN, Callosal Projection Neurons; ScPN, Subcerebral Projection Neurons; CThPN, CorticoThalamic Projection Neurons.

*Related to main Figure 4.*

**Figure S10**

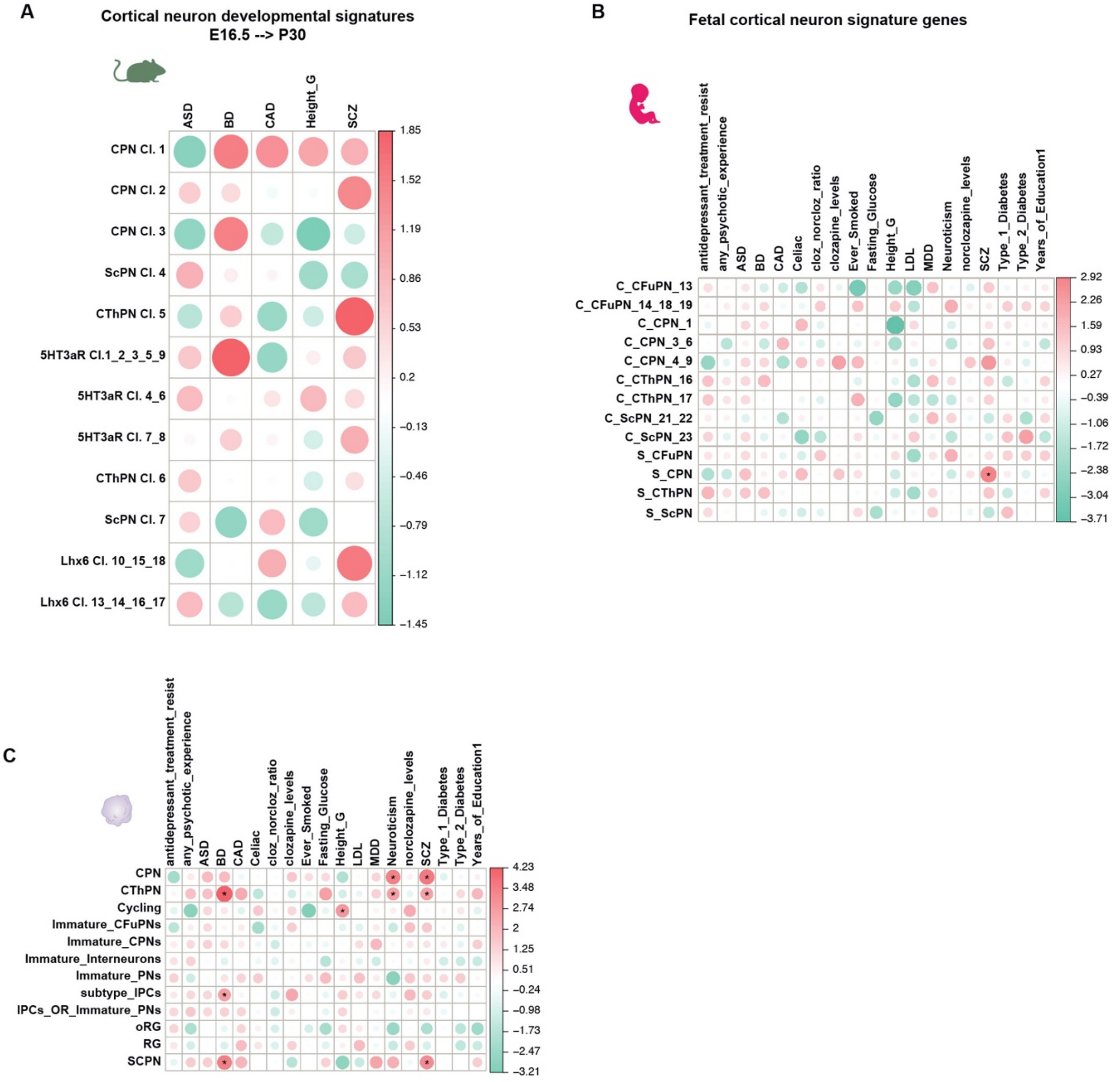

A
Cortical neuron developmental signatures
E16.5 --> P30
ASD
BD
CAD
Height_G
SCZ
CPN Cl. 1
CPN Cl. 2
CPN Cl. 3
ScPN Cl. 4
CThPN Cl. 5
5HT3aR Cl.1_2_3_5_9
5HT3aR Cl. 4_6
5HT3aR Cl. 7_8
CThPN Cl. 6
ScPN Cl. 7
Lhx6 Cl. 10_15_18
Lhx6 Cl. 13_14_16_17
1.85
1.52
1.19
0.86
0.53
0.2
−0.13
−0.46
−0.79
−1.12
−1.45
B
Fetal cortical neuron signature genes
antidepressant_treatment_resist
any_psychotic_experience
ASD
BD
CAD
Celiac
cloz_norcloz_ratio
clozapine_levels
Ever_Smoked
Fasting_Glucose
Height_G
LDL
MDD
Neuroticism
norclozapine_levels
SCZ
Type_1_Diabetes
Type_2_Diabetes
Years_of_Education1
C_CFuPN_13
C_CFuPN_14_18_19
C_CPN_1
C_CPN_3_6
C_CPN_4_9
C_CThPN_16
C_CThPN_17
C_ScPN_21_22
C_ScPN_23
S_CFuPN
S_CPN
S_CThPN
S_ScPN
2.92
2.26
1.59
0.93
0.27
−0.39
−1.06
−1.72
−2.38
−3.04
−3.71
C
antidepressant_treatment_resist
any_psychotic_experience
ASD
BD
CAD
Celiac
cloz_norcloz_ratio
clozapine_levels
Ever_Smoked
Fasting_Glucose
Height_G
LDL
MDD
Neuroticism
norclozapine_levels
SCZ
Type_1_Diabetes
Type_2_Diabetes
Years_of_Education1
CPN
CThPN
Cycling
Immature_CFuPNs
Immature_CPNs
Immature_Interneurons
Immature_PNs
subtype_IPCs
IPCs_OR_Immature_PNs
oRG
RG
SCPN
4.23
3.48
2.74
2
1.25
0.51
−0.24
−0.98
−1.73
−2.47
−3.21

**Figure S10: LD regression score for GWAS hits for neurodevelopmental and neuropsychiatric disorders in subtype-specific signatures of human and mouse cortical classes by cluster.** **A**) Genetic correlations (estimated by LD Score Regression) of a wide array of diseases did not show association with any mouse PN and IN signature set, even when each population is clustered separately. **B**) Genetic correlation with human fetal clusters confirmed significant association with SCZ and CPN subtype. **C**) Genetic correlation with human 3mo cortical organoids (Table S10) confirmed significant association with SCZ and CPN subtype, in addition to CThPNs and ScPNs, and significant enrichment for neuroticism in CPN and CThPN, as well as BD in ScPNs. *represents significant genetic correlation. A full list of the sum statistics of the traits are reported in Table S10.

Abbreviations: Cl, cluster; S, subtype; CPN, Callosal Projection Neurons; ScPN, Subcerebral Projection Neurons; CThPN, CorticoThalamic Projection Neurons; CFuPNs, Corticofugal Projection Neurons; IPCs, intermediate progenitor cells: PNs, Projection Neurons; oRG, outre Radial Glia; RG, radial glia.

*Related to main Figure 5.*
